# Population genetics of the endangered narrowly endemic Island Marble butterfly (*Euchloe ausonides insulanus*)

**DOI:** 10.1101/2025.05.08.652852

**Authors:** Kara S. Jones, Aaron W. Aunins, Colleen C. Young, Robin L. Johnson, Cheryl L. Morrison

## Abstract

The Island Marble butterfly (*Euchloe ausonides insulanus*) is an endangered species endemic to the San Juan Islands off the coast of Washington State, United States, and British Columbia, Canada. The species was thought to be extinct for ∼90 years before it was rediscovered at American Camp, San Juan Island National Historical Park in 1998. Here, we report the results of the first population genetic analyses for *insulanus*, using DNA collected non-invasively from individuals in the last known stronghold for the species. We used DNA extracted from meconium, larval exuviae, and natural mortalities to generate and test thirteen new microsatellite markers to estimate genetic diversity, population structure, and kinship. We assembled and annotated mitochondrial genomes, which were used alongside museum specimens of *insulanus* collected ∼100 years ago from Vancouver Island, and other members of the *E. ausonides* species complex, to infer the evolutionary history of the species. The results indicated that *insulanus* experiences low heterozygosity, a small effective population size (N_e_), and low allelic diversity. High levels of inbreeding were found in some individuals, but inbreeding was uneven across the population. No population structure or partitioning of genetic variation by host plant was detected. The mitogenomes of extant *insulanus* were all identical and modern samples showed a loss of allelic diversity compared to *insulanus* from museums. Extant *insulanus* formed a clade with museum specimens and we identified multiple putatively diagnostic alleles to differentiate *insulanus* from other subspecies. Based on these results, we outline considerations for species management and genetic monitoring.

## Introduction

The Island Marble butterfly (*Euchloe ausonides insulanus*), hereafter referred to as *insulanus*, is a small butterfly in the family Pieridae that is endemic to the San Juan Islands off the coast of Washington State, United States and British Columbia, Canada. The last specimen was collected from Gabriola Island, British Columbia, in 1908, and was believed to be extinct for ∼90 years before the butterfly was rediscovered at American Camp, San Juan Island National Historical Park (SAJH NHP) in 1998 (Shepard, 2000). Extensive survey efforts identified five areas of occupied habitat across two islands (San Juan Island and Lopez Island), but subsequent habitat loss has led to widespread declines in the distribution of the species (Lambert, 2011). By 2012, the species distribution had contracted to a single population mainly within the boundaries of American Camp. Due to the declining status of *insulanus*, an emergency captive-rearing effort to sustain the last remaining population began in 2013. In 2020, *insulanus* was listed as an endangered species (U.S. Fish and Wildlife Service 2020; 85 FR 26786) under the Endangered Species Act (ESA 1973, as amended), and 812 acres on the south end of San Juan Island were designated critical habitat (50 CFR 17.95(i)).

Several life history traits may contribute to the current precarity of *insulanus*. Based on Lambert’s (2011) detailed natural history of the species, oviposition occurs on three larval host plants in the mustard family (Brassicaceae), each of which primarily occurs in discrete habitats: *Brassica rapa* (grasslands), *Sisymbrium altissimum* (sand dunes), and *Lepidium virginicum* var. *menziesii* (tidal zones). Larvae develop through five instars, taking ∼38 days to progress from egg to pupa, after which pre-pupal larvae “wander” up to four meters to a different species of plant to pupate for 10-11 months. Over 50% of eggs are lost prior to hatching, approximately half from predation on host plants by deer. Larvae are vulnerable to death from starvation, particularly when feeding on host plants growing in marginal habitat, where the host plants may senesce before larvae reach the pre-pupal stage. Furthermore, *insulanus* is univoltine (i.e., has non-overlapping generations), making the species particularly vulnerable to the effects of storms and localized climate conditions during the long winter diapause as pupae (McDermott Long et al., 2017). The larval host plants *S. altissimum* and *L. virginicum* var. *menziesii*, are located primarily in sand dunes and tidal habitats, respectively, which increases the vulnerability of host plants to wind, wave action, and other disturbances. This can lead to fewer oviposition sites, higher larval mortality, or higher pupal mortality, depending on the time of year (Lambert, 2011). Hence the combined stressors of loss and degradation of habitat, storm surges, direct and incidental predation, and limited larval host plant availability, may have contributed to the extirpation of *insulanus* from Vancouver, Gabriola, and Lopez Islands and contraction of the species’ range on San Juan Island (Shepard 2000; U.S. Fish and Wildlife Service 2023a).

Previous reports examining the status of *insulanus* identified several information gaps that could be addressed by genetic data, including neutral genetic diversity, effective population size, and population structure (U.S. Fish and Wildlife Service 2023a, 2023b, 2023c). However, there are several challenges and limitations to consider when generating genetic markers for an imperiled species. For example, non-invasive sampling techniques are usually required to minimize harm to the organism being studied when collecting DNA samples. In the case of *insulanus*, DNA collection is mostly limited to lower-quality sources, such as frass or exuviae shed between larval instars, potentially resulting in fewer loci than data sets obtained by extracting DNA from fresh tissue samples (Schultz et al., 2022). Non-invasively sampled DNA can also lead to an increased number of genotyping errors due to allele dropout from partially degraded templates (Zhan et al., 2010), errors introduced from an increased number of amplification cycles (Arantes et al., 2023), and sequence contamination with DNA from non-target organisms and the surrounding environment (Balmori-de la Puente et al., 2023; Sittenthaler et al., 2023). Moreover, non-invasively collected samples are usually collected opportunistically, and therefore the individuals included in genetic analyses may not be representative of the population, which can bias results (Waits and Leberg, 2000). Hence, genetic studies of threatened and endangered species require careful consideration at each step to correct errors and reduce bias introduced through suboptimal sampling procedures.

In this study, we assessed the quality of DNA that could be recovered from available non-invasive sampling templates (e.g., meconium, chrysalides, exuviae, and frass) collected from *insulanus* being housed at the SAJH NHP Captive Rearing Facility. We developed a suite of microsatellite markers, which we assessed for information content and biases that may have been introduced by sampling constraints, before using the markers to estimate genetic diversity, levels of inbreeding, and effective population size. We used kinship analyses to identify cryptic relatedness and create a reduced- relatedness data set to explore whether the population was structured by larval host plant use. We also used mitochondrial genomes to explore the evolutionary history of the species in relation to other subspecies in the complex. Finally, we place the results in the broader conservation planning initiative for *insulanus*, particularly regarding captive rearing and translocation efforts.

## Materials and Methods

### Sample collection

Samples of *insulanus* used for this study were collected non-lethally by researchers at the SAJH NHP Captive Rearing Facility from 2013 through 2019, prior to *insulanus’* listing as an endangered species. A variety of sample types were collected for DNA, including tissue from natural mortalities (e.g., eggs, larvae), and discarded solids (e.g., chrysalides, exuviae, frass) and liquids (e.g., meconium) (Table 1). Tissues and solid samples were collected using gloved hands and clean forceps, and samples were submerged in lysis buffer or 95% ethanol in 2 μl sterile screwcap tubes. Liquid meconium samples were spotted on FTA cards (Cytiva, Wilmington, Delaware), and the cards were stored in sealable plastic storage bags with desiccant. Samples were shipped with an ice pack to USGS EESC-LRL for further processing. Sample metadata are listed in Supplementary Table S1.

**Table 1.**
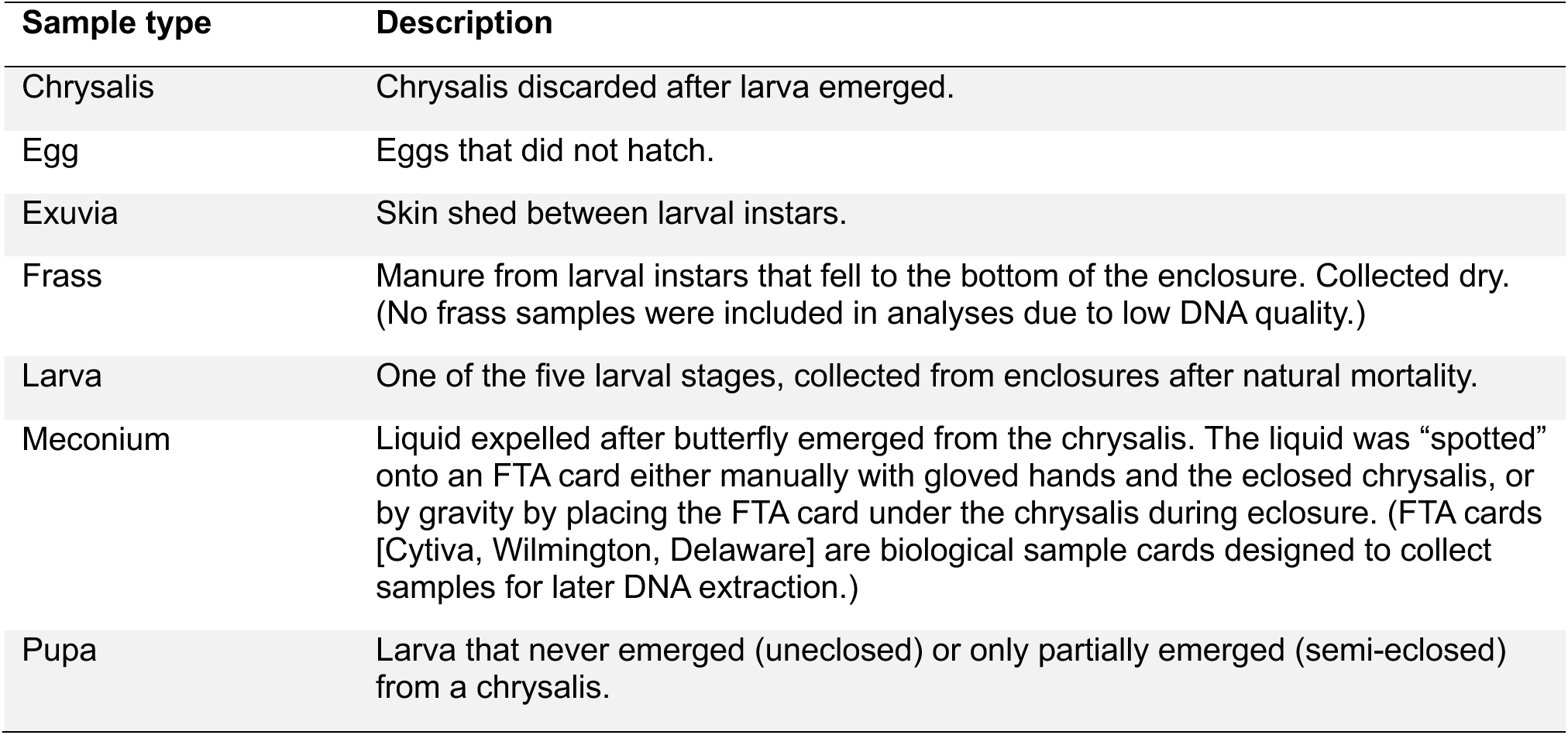
Types of non-invasive samples collected for DNA. All samples were collected either dry, in 95% ethanol, or in Qiagen DNEasy Cell Lysis Buffer (Germantown, Maryland), except where noted.

### Microsatellite marker development and genotyping

DNA was extracted from the meconium of a captive *insulanus* butterfly (Eai-S43) using the Qiagen DNeasy Blood and Tissue Kit (Germantown, Maryland) following the manufacturer’s protocol. A genomic DNA library was prepared for Illumina sequencing using the TruSeq Nano kit (San Diego, California), and 2x150 bp paired-end (PE) and sequenced on a NextSeq500 (San Diego, California) at the USGS EESC-LRL. The resulting sequences were quality trimmed using Qiagen CLC Genomics Workbench (CLCGW) v9.5.2 (Qiagen, 2016) using a quality limit value of 0.05 (corresponding to a Phred score of ∼13), and a minimum sequence length of 15 bp. Cleaned sequences were then *de novo* assembled (i.e., assembled without a reference) into contigs using CLCGW with the following parameters: automatic bubble and word size, minimum contig length of 200, scaffold assembly enabled, and auto-detection of paired distances.

We screened contigs for di-, tri-, tetra-, penta-, and hexanucleotide microsatellite repeat motifs with a minimum repeat length of five using the program QDD v3 (Meglécz et al., 2010). Loci were aligned to the GenBank Nucleotide database (Benson et al., 2017) using BLAST (Camacho et al., 2009) and discarded if they could be assigned to a probable contaminant (e.g., fungi or bacteria). Primer sequences for the microsatellite loci were identified using Primer 3 within QDD, with forward primers modified to contain a universal M13 tail to enable the incorporation of a fluorescent dye label during PCR for fragment analysis.

Seventy-four microsatellite primer pairs were chosen for testing. Microsatellite DNA amplification consisted of 5-40 ng of genomic DNA, 5X PCR buffer (10 mM Tris-HCl, pH 8.3, 50 mM KCl), 2 mM MgCl_2_, 0.25 mM dNTPs, 0.25 µM forward or long M13 tagged primer, 0.5 µM reverse or short primer, 0.1 µM M13 fluorescently labeled primer, 0.13 U Bovine Serum Albumin (BSA) and 0.1 U Taq DNA polymerase (Promega, Madison, WI, USA) in a final reaction volume of 15 µl. Each locus was amplified separately on a T100 Thermal Cycler (Bio-Rad, Hercules, California) using the following conditions: initial denaturing at 94°C for 15 min, 29 cycles of 94° C for 1 min, 60°C for 45 sec, 72 ° C for 45 sec, followed by 5 cycles of 94° C for 1 min, 53° C for 45 sec, 72 ° C for 45 sec and a final extension at 72 ° C for 10 min. Fragment length polymorphism was assessed using a ThermoFisher ABI 3130 Capillary Sequencer (Waltham, Massachusetts) and ThermoFisher’s GeneMapper v4.1 software (Applied Biosystems, 2009).

Thirteen loci amplified consistently across samples and were polymorphic. These primer sets were ordered with ABI fluorescent dyes and without M13 tails and were arranged into multiplex PCR reactions (one duplex, one triplex, two quadruplexes; Table S2) using Multiplex Manager (Holleley and Geerts 2009). These new multiplex PCRs were used to genotype additional samples, most of which were from meconium preserved on FTA cards, following the same scoring on the ABI 3130.

Summary statistics for each microsatellite locus, including the number of alleles (*N*_a_), effective number of alleles (*A*_e_), observed heterozygosity (*H*_o_), expected heterozygosity (*H*_e_), fixation index (*F*), and tests for conformance to Hardy-Weinberg equilibrium (HWE), were calculated in GenAlEx (Peakall and Smouse, 2012).

### Microsatellite kinship, inbreeding, and relatedness estimates

Population genetic analyses can be biased when using relatively few markers and/or markers with low information content. We therefore used the *pic_calc* function from the PopGenUtils package (Tourvas, 2024) in R 4.4.1 (R Core Team, 2024), to calculate the polymorphic information content (PIC) of each locus. We also used the *pid_calc* function to calculate the probability of identity for two unrelated (P_(ID)_) or two related (P_(ID)sib_) individuals. PIC measures how much variation occurs at a locus within a population, providing an estimate of whether a marker can detect genetic differences between individuals, and P_(ID)_/P_(ID)sib_ estimates the probability that two individuals will have the same genotype (Waits et al., 2001; Serrote et al., 2020). One locus (Locus-71) was dropped from all subsequent analyses because of low PIC and high P_(ID)_/P_(ID)sib_.

We estimated sibship among individuals, effective population size (N_e_), and allelic error rates using COLONY 2.0.7 (Wang, 2009, 2018; Jones and Wang, 2010). Since the life cycle of *insulanus* lasts for one year and adults die after breeding, samples were grouped into three offspring-parent cohorts. Genotypes from 2017, 2018, and 2019 were each set as offspring and analyzed with candidate parent genotypes from the previous year (e.g., 2017 samples as offspring were analyzed with 2016 samples as parents). The sex of the samples was unknown, so the same set of genotypes was used for both maternal and paternal candidates. To ensure consistency across results, we ran each cohort with the full likelihood method using the “very high” precision setting and a “very long” run length (i.e., the most precise and longest lengths possible for the program). Each cohort was run for 10 replicates using the following settings: dioecious species (2), inbreeding allowed (1), polygamous males (1) and females (1), no clone inference (0), full sibship size scaling (1), no sibship size prior (0), unknown population allele frequency (0), allele frequency updates (1). A starting error rate of 1% was used for estimating allelic dropout and false alleles, and the probability that an actual mother or father was included in candidate parent genotypes was set at 1%.

We used EMIBD9 1.1.0.0 to estimate inbreeding coefficients (*F*; i.e., the probability that two alleles at a specific locus in an individual are identical by descent) for each individual and to calculate pairwise relatedness (*r*) for all individuals across all years (Wang, 2022). To determine whether significant variance in inbreeding was present among individuals, we calculated g_2_ with the *g2_microsats* function in the inbreedR package, using 10,000 permutations to obtain the p-value and 10,000 bootstraps to calculate 95% confidence intervals (Stoffel et al., 2016). To determine whether the *insulanus* population was being structured by host plant use, we evaluated the difference in mean pairwise relatedness between individuals found on the same host plants versus different host plants with two-sided t-tests using the Infer package in R (Couch et al., 2021).

Since the inclusion of highly related individuals can lead to clustering by kinship and obfuscation of other patterns in the data (Falush et al., 2007), we created a reduced-relatedness data set by removing all but one individual from each full sibship group with ≥0.9 inclusive probability assigned by COLONY. We performed a principal components analysis (PCA) (Patterson et al., 2006) on this reduced data set (N = 193) with the *dudi.pca* function of the adegenet 2.1.10 (Jombart, 2008) package in R to explore whether there was structure in the data that could be attributable to differential host plant use. Additionally, we ran Structure 2.3.4 (Pritchard et al., 2000) for population clusters (K) of 2, 3, and 4 using a burn-in of 100,000 steps and 500,000 MCMC steps. The MCMC chain was considered to have convergence when the likelihood values had minimal variance between steps. Individuals were split into groups by year, and each year and value of K was run ten times independently. Data was summed across runs using Clumpak (Kopelman et al., 2015). The PCA and Structure results were visualized in R using tools in the tidyverse set of packages (Wickham et al., 2019).

### Mitogenome sequencing, assembly, and annotation

We compared shotgun sequencing with Long Range PCR (LR-PCR) to assess which method was more effective at recovering mitogenomes with low template quantity and/or quality (Ponce and Micol, 1992). For shotgun sequencing, seven *insulanus* samples and two samples from other *Euchloe ausonides* subspecies were prepared for shotgun genomic sequencing on an Illumina MiSeq (San Diego, California) using the New England Biolabs NEBNext Ultra II DNA Library Prep Kit for Illumina (Ipswich, Massachusetts). Samples were dual indexed with combinatorial indices and paired-end sequenced at EESC-LRL across two 500-cycle MiSeq cartridges. For LR-PCR, we used three *insulanus* and one *E. a. subsp.* from the same pool of individuals as above. LR-PCR primers were adapted from Park et al. (2012) to better match *Euchloe* sequences in GenBank (Table S3). Each mitogenomic fragment was amplified in a 50 µl PCR reaction consisting of 0.5 µl TaKaRa LA Taq (San Jose, California; 5U/µl stock concentration), 5 µl 10X LA PCR Buffer, 5 µl 25 nM MgCl_2_, 8 µl dNTP mixture (2.5 mM each stock concentration), 1 µl forward and 1 µl reverse primers (each at 10 µM stock concentration) 2 µl DNA template, and 27.5 µl PCR grade water. Thermal cycling parameters for each of the fragments were 94°C for 1 min, followed by 35 cycles of 98°C for 10 s, relevant annealing temperature (AT; degrees C) from Table S3 for 15 min, 72°C for 30 s, and a final extension at 72°C for 10 minutes. These fragments were purified and pooled in equimolar concentrations and then subjected to the same library preparation protocol as the shotgun libraries for Illumina sequencing (as described in the section *Microsatellite marker development and genotyping*).

Reads from WGS sequencing and LR-PCR were *de novo* assembled with MitoZ 3.6 (Meng et al., 2019) using Megahit (Li et al., 2015) for the assembler, and substrings of length k (k-mers) of 59, 79, 99, 119, 141. For comparison, a subset of individuals (EaC3, 17SI21, Eai-163) was also assembled with MitoFinder 1.4.1 (Allio et al., 2020), using the annotated *Anthocharis bambusarum* mitogenome (NCBI RefSeq: NC_025274) as reference (Ebdon et al., 2022). The resulting mitogenomes were annotated with MITOS2 2.1.9 and mitochondrial tRNA finder (MiTFi) on the public server Galaxy (usegalaxy.org), using a genetic code = 5 for translation and RefSeq89 Metazoa as a reference (Al Arab et al., 2017; Afgan et al., 2018; Donath et al., 2019). Additional tRNA annotation was performed using tRNAscan-SE 2.0 (Chan et al., 2021). Annotations were compared with existing mitochondrial genomes to ensure they had a similar length and structure. Protein coding genes were also verified by checking for the presence of start and stop codons, absence of stop codons within the gene, and similarity to genes found in other butterfly mitogenomes found in GenBank.

### Mitogenome phylogeny

To determine how the *insulanus* mitogenome differed from those of other subspecies, we obtained whole mitogenome sequences that had been assembled for an unrelated *Euchloe* project (Zhang et al., University of Texas Southwestern Medical Center, unpublished data, 2023). The final data set had mitogenomes from six individuals of *E. a. ausonides*, six *E. a. coloradensis*, three *E. a. mayi*, four *E. a. palaeoreios*, eight *E. a. transmontana*, and five additional individuals of *insulanus* derived from museum specimens collected from Vancouver Island circa 1898-1904 (Table S4). All assemblies were rotated to the same starting position using Cyclic DNA Sequence Aligner (Fernandes et al., 2009), aligned using MUSCLE 3.8.31 (Edgar, 2004) and visually assessed for consistency in AliView 1.28 (Larsson, 2014). Single nucleotide polymorphisms were extracted from the aligned mitogenomes using SNP-sites (Page et al., 2016).

With the addition of a sequence from *E. lotta* to be used as an outgroup, we used the aligned mitogenomes to infer a phylogeny with IQTree2 2.2.2.6 (Minh et al., 2020). Mitochondrial sequences were partitioned by feature, with coding regions additionally partitioned by codon position and model selection chosen with the MFP+MERGE setting (Chernomor et al., 2016). Node support was assessed with ultrafast bootstraps and site concordance factors (sCF), both calculated using 1,000 iterations (Hoang et al., 2018; Mo et al., 2023).

## Results

### Microsatellites

The sequencing run of Eai-S43 generated 94,216,390 raw read pairs, of which 93,522,078 read pairs remained after quality and length trimming. *De novo* assembly resulted in 345,212 contigs with an average length of 395 bp (N50 = 412 bp, min size = 200 bp, max size = 15,385 bp).

QDD identified 7,370 candidate microsatellite sequences from the assembled contigs of Eai-S43, of which 74 were chosen for testing. Nineteen of the 74 loci chosen for testing yielded two or more alleles, indicating they were potentially polymorphic for a larger number of alleles. Thirteen of these loci were optimized for multiplex PCR and genotyping (Table S2). DNA from the whole desiccated larvae provided the highest quality and quantity of DNA, followed by meconium preserved on FTA cards. Frass was found to be an unsuitable template due to poor DNA quality and quantity. Overall, 251 samples that were genotyped were distributed unevenly across years, with 14 individuals from 2016, 56 from 2017, 37 from 2018, and 44 from 2019 (Table S1).

The number of alleles per locus ranged from two to nine, but was low overall, with most loci only having two alleles (Table S5). The locus with the highest number of alleles was Locus-24 with seven. Testing for conformance to HWE indicated that 7 of the 13 loci in the 2018 cohort were out of HWE after Bonferroni correction, and observed heterozygosity (*H_o_*) was lower than expected for all individuals. The average *H*_o_ across loci was 0.41 for the 2016 samples, 0.42 for the 2017 samples, and 0.40 for the 2018 samples indicating a relatively low level of heterozygosity in all three populations.

Polymorphic information content (PIC) ranged from 0.06-0.67, P_(ID)_ from 0.13-0.87, P_(ID)sib_ from 0.43-0.93 (Table S6). Locus L-71 scored particularly poorly across all metrics, having the lowest PIC and highest P_(ID)_/P_(ID)sib_ and was therefore removed from subsequent analyses. After removal of L-71, H_o_ increased for 2016 and 2018 cohorts, and decreased for 2017 and 2019, whereas fixation index (F_IS_) estimates increased for 2017 and 2018 (Table 2).

**Table 2.** Number of alleles (A), effective number of alleles (A_e_) observed heterozygosity (H_o_), expected heterozygosity, (H_e_) and fixation index (F_IS_) for each year using 12 microsatellite loci (after L-71 was removed due to low information content).

| | N | A | $A_e$ | $H_o$ | $H_e$ | $F_{IS}$ |
| --- | --- | --- | --- | --- | --- | --- |
| **2016** | 14 | 2.25 ± 0.131 | 1.92 ± 0.147 | 0.37 ± 0.053 | 0.44 ± 0.044 | 0.133 ± 0.096 |
| **2017** | 56 | 2.42 ± 0.336 | 1.82 ± 0.097 | 0.38 ± 0.039 | 0.43 ± 0.032 | 0.106 ± 0.061 |
| **2018** | 137 | 2.67 ± 0.414 | 1.86 ± 0.133 | 0.33 ± 0.049 | 0.43 ± 0.042 | 0.218 ± 0.076 |
| **2019** | 44 | 2.33 ± 0.188 | 1.86 ± 0.108 | 0.45 ± 0.045 | 0.44 ± 0.035 | -0.03 ± 0.054 |
| **Mean** | 62.75 | 2.42 ± 0.142 | 1.86 ± 0.06 | 0.39 ± 0.023 | 0.44 ± 0.019 | 0.107 ± 0.038 |

Using COLONY, individuals were assigned to 20 full sibship groups in the 2017 cohort, 28 in 2018, and 15 in 2019 (Table S7). Across all years, there were 26 full sibship groups with ≥0.9 inclusive probability (total membership = 78). The fixation index (F_IS_) per cohort was estimated at 0.09-0.17 and N_e_ was 31-47 (Table 3). Estimated false allele and allele dropout rates varied widely (0-50%) by year and marker (Figure S1), averaging 6.6% and 5%, respectively.

**Table 3.**
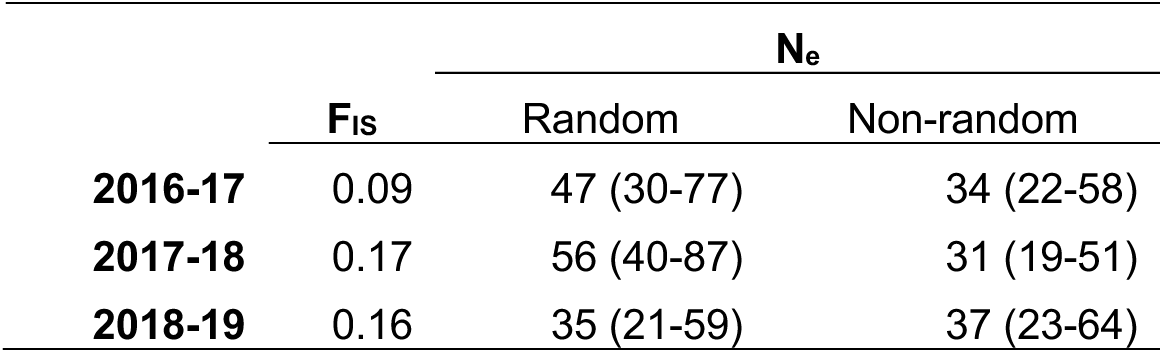
Estimates from COLONY of inbreeding (i.e., the fixation index, F_IS_) and effective population size (N_e_) and 95% confidence interval (in parentheses) for three sibling cohorts. N_e_ was calculated either assuming random or non-random mating.

Inbreeding coefficients varied widely across individuals (range of *F* = 0.006-0.903; Figure S2), and there was significant variance in inbreeding among individuals (g_2_ = 0.06 ± 0.018, p = 0.0001).

Mean pairwise relatedness between individuals was similar across years (*r* = 0.13-0.15) but there were more highly related pairs (*r* > 0.5; roughly full sibling or higher) in 2018 than other years (25% vs 2- 5%), which could have impacted within-year estimates of H_o_ and F_IS_, as well as clustering patterns in population structure analyses. Overall, individuals collected in the same year had significantly higher *r* values compared to those collected from different years (*t* = 23.6, *df* = 23,005, p = 4.7 x 10^-122^). There was no significant difference in relatedness between individuals from the same or different host plants (*t* = 1.39, *df* = 13,917, p = 0.164).

The first two axes of the PCA captured 27.5% of the variance. No combination of axes showed discernable clustering by year or host plant (Figure 1). Structure analyses showed no clear clustering by host plant at any values of K (Figure 2 and S3). Individuals in three years (2016, 2017, and 2019) showed equal proportions of ancestry assigned to each cluster, while 2018 showed some variability in cluster assignment among individuals, possibly due to the higher relatedness among individuals from that year class.

**Fig. 1.**
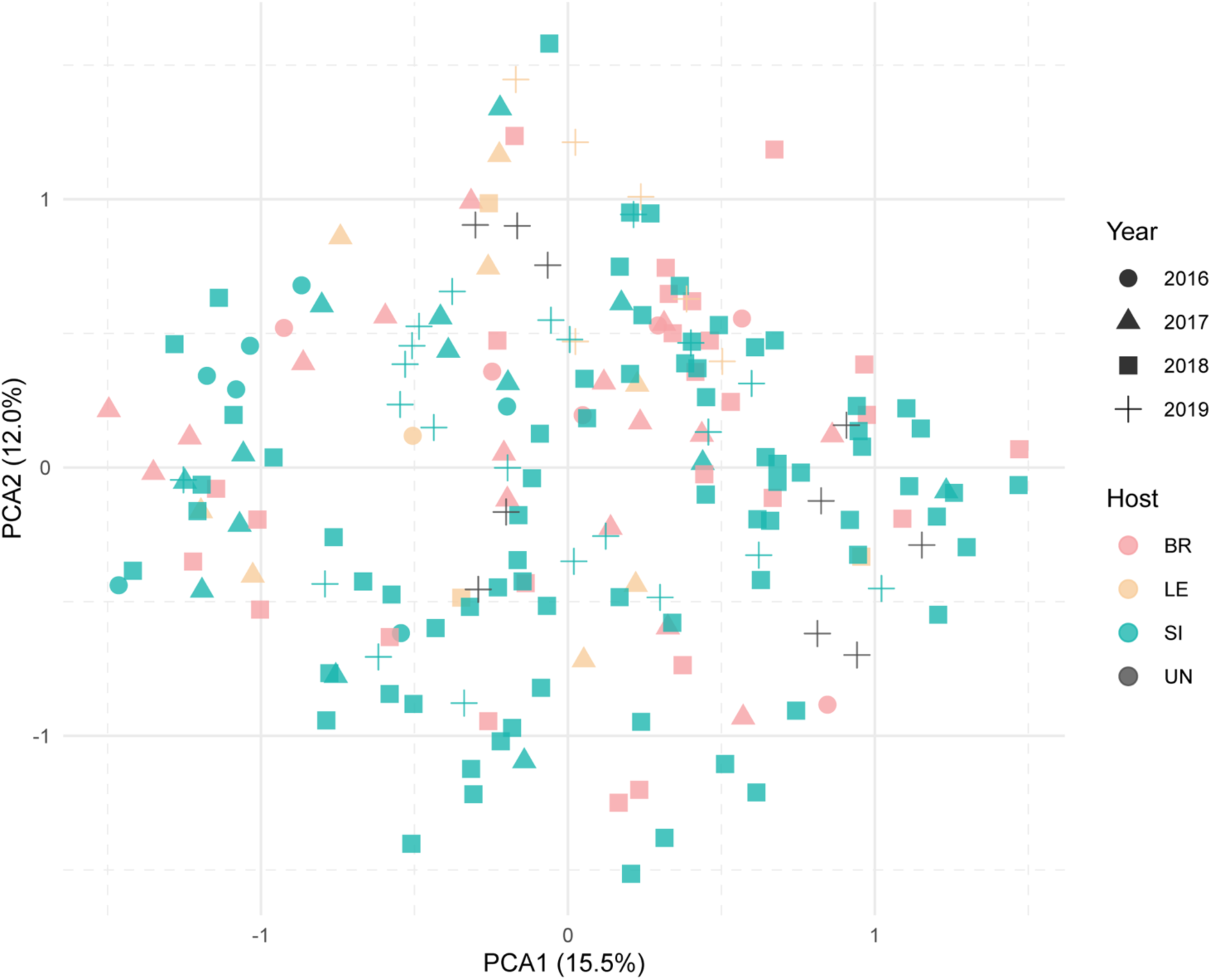
Principal Component Analysis (PCA) using non-related individuals from all years, colored by larval host plant (BR = *Brassica*, LE = *Lepidium*, SI = *Sisymbrium*, UN = larval host plant not recorded). Percent variance explained by each axis shown in parentheses. There was no clear signal of structure by year or host plant.

**Fig. 2.**
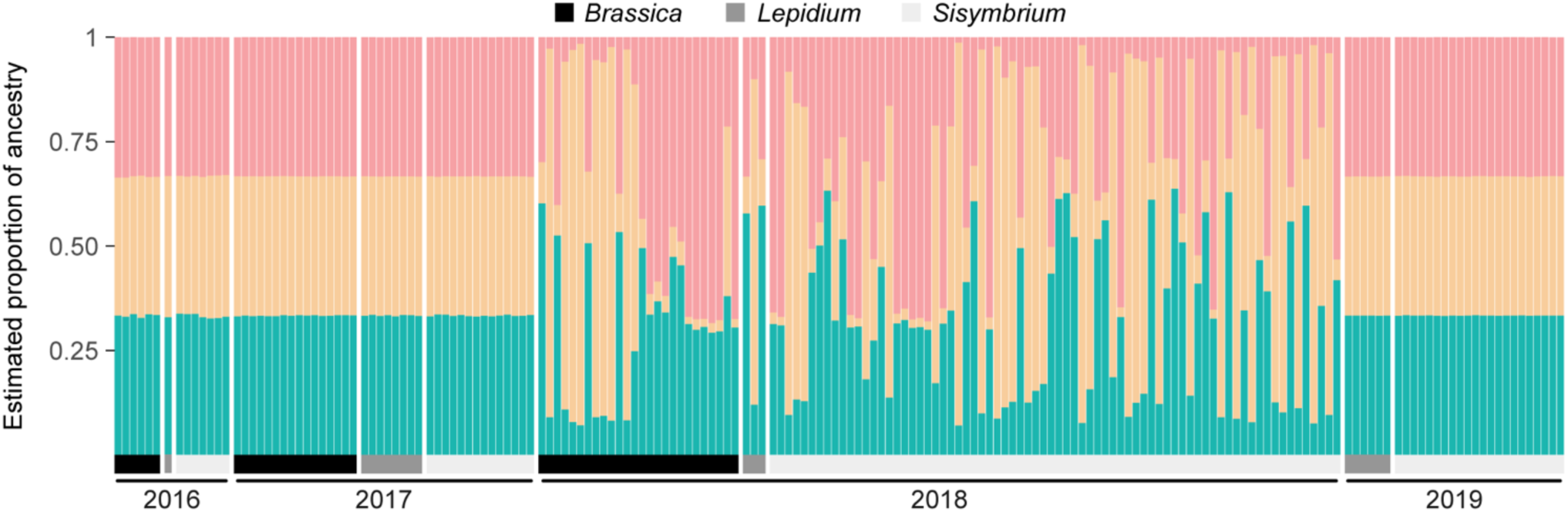
Comparison of estimated ancestry from Structure for population clusters (K) = 3. Each year was run through Structure separately using a reduced-relatedness data set created by removing all but one individual from each of the full sibship clusters (≥0.9 inclusive probability) output from COLONY. Individuals are grouped by larval host plant genus within each year and colors indicate estimated proportion of estimated ancestry to each cluster.

### Mitogenomes

We assembled nearly complete circular mitogenomes for all mitogenomic samples except for Eai-162 and EaC8, which only had partial sequences for several genes and were therefore discarded. Mean read depth along the mitogenomes was 63-49,449 reads for shotgun sequencing and 10,833- 23,195 reads for LR PCR (Table S8). The only areas of the mitogenome that showed variation among *insulanus* assemblies were indels found in the AT-rich control region. However, there were also spikes in read depth in this region which exceeded the coverage across the rest of the mitogenome by >10x, indicating that mapping quality in the control region was likely poor due to repeats. We trimmed 2 bp indels at positions 13,426-13,427 and 13,171-13,172 (relative to the start of the Cytochrome c oxidase subunit 1 gene) to remove possible erroneous assemblies but the control region may still contain errors and/or be incomplete. All *insulanus* mitogenomes were 15,193 bp long, contained the expected 13 protein-coding genes, 22 tRNAs, 2 rRNAs, D-loop, and replication origin (OH), and had identical sequences after trimming (Figure S4).

We found nine SNPs that are putatively diagnostic for extant *insulanus*, three of which were not found in the museum specimens (Table 4). One SNP was unique to museum *insulanus* but not found in the extant samples, whereas three were variable in museum specimens but are now fixed in extant *insulanus*, potentially representing a loss of historical diversity. There was high node support (bootstrap [B] = 98%, site concordance factor [sCF] = 98%) for extant and museum *insulanus* forming a monophyletic clade (Figure 3). There was similarly high support (B = 98%, sCF = 98%) for the *insulanus* clade being placed as sister to *E. a. mayi*, rather than the geographically adjacent mainland subspecies *E. a. transmontana*. However, we note that relationships among subspecies were generally poorly resolved, which may reflect a lack of phylogenetic information in the mitogenomes and/or a mismatch between genetic divergence and current taxonomy.

**Fig. 3.**
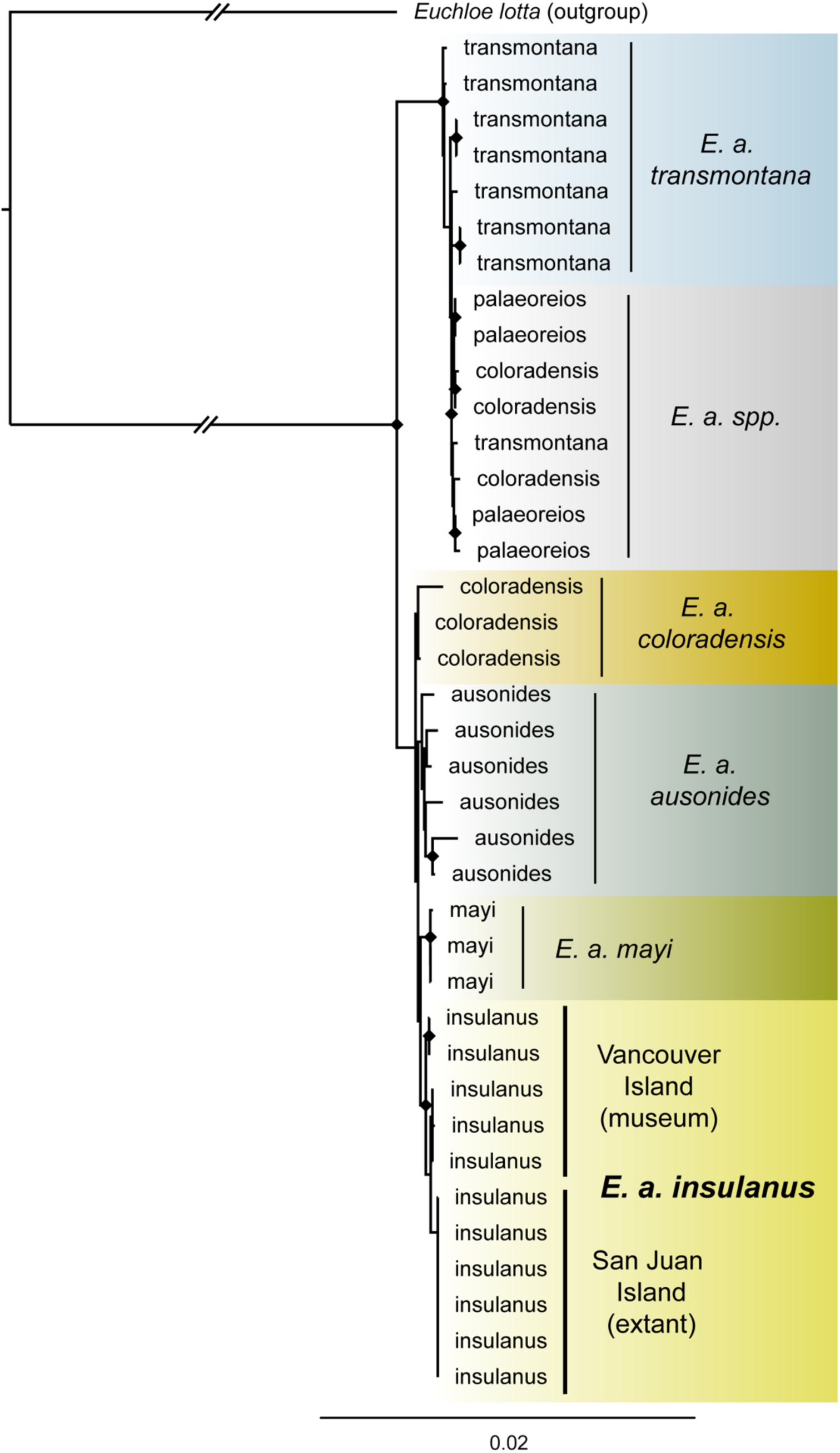
Mitogenome phylogeny for *Euchloe ausinodes* subspecies, including *insulanus*, using extant samples from San Juan Island and museum specimens from Vancouver Island. Diamonds represent node support with site concordance factor (sCF) ≥ 0.9. All insulanus specimens (extant and museum) formed a monophyletic clade with high node support (bootstrap = 98%, sCF = 98%), placed sister to *E. a. mayi*. The mitogenome for *insulanus* is available under GenBank accession PQ287242.1. At the time of publication, additional sequences used in the mitogenome phylogeny from the University of Texas Southwestern Medical Center were not available for publication.

**Table 4.** Mitochondrial single nucleotide polymorphisms (SNPs) found in *Euchloe ausonides insulanus* compared to other *E. ausonides* subspecies. Positions are relative to start of COX1 gene. All SNPs listed are potentially diagnostic for extant *E. a. insulanus*, except for position 6161, which was only found in museum *insulanus* specimens and is therefore likely to have been lost. Reference alleles are those found in other *E. ausonides* subspecies included in this study. * Potentially unique to *insulanus* on San Juan Island; ** Potentially diagnostic for extant *insulanus* population; † Allele likely lost in extant *insulanus* population. Mitogenome for insulanus available under GenBank accession PQ287242.1. At the time of publication, additional sequences used in the mitogenome phylogeny from the University of Texas Southwestern Medical Center were not available for publication.

| Position | Reference | *E. a. insulanus* |
| --- | --- | --- |
| 702 | A | G * |
| 1201 | C | T ** |
| 1685 | G | A * |
| 6161 | T | C † |
| 6780 | T | C ** |
| 7858 | A/T | C ** |
| 8516 | A | G ** |
| 10084 | T | C ** |
| 14167 | C | T ** |
| 14347 | T | C ** |

## Discussion

This study offers a first look into the genetic diversity of the endangered butterfly species *Euchloe ausonides insulanus*. We generated microsatellite markers from DNA collected non-invasively from exuviae, meconium, and other sources, showing the potential of monitoring genetic diversity in this species without relying on lethal sampling techniques or incidental mortalities. Additionally, the annotated mitochondrial genome presented here for *insulanus* sheds light on the genetic diversity within the species and its relationship to other members of the *E. ausonides* species complex. Our work also provides putatively diagnosable genetic markers to distinguish *insulanus* from other subspecies in the complex.

Our results indicate that *insulanus* has relatively low heterozygosity, a small effective population size (N_e_), and low allelic diversity. These factors limit the available genetic variation the species has to draw from and reduce the likelihood that advantageous mutations will arise in the future, both of which decrease the species’ adaptive potential (Hoffmann et al., 2017; Rousselle et al., 2020). The number of alleles per locus and observed heterozygosity estimated with microsatellites was lower than those found in most previous butterfly studies (as reviewed in Heffernan et al. 2024), and on par with some declining species, such as the endangered Poweshiek skipperling (*Oarisma poweshiek*) (Saarinen et al., 2016).

Furthermore, an N_e_ of 50-500 may be necessary to avoid extinction (Frankham et al., 2014; Pérez-Pereira et al., 2022), and our estimates for *insulanus* were well below that minimum. The lack of population structure and low genetic diversity also indicate that *insulanus* likely exists as a single population isolated on one island with no connectivity to other subspecies that could provide additional genetic diversity through gene flow. These circumstances may restrict *insulanus’* access to the genetic diversity needed to assemble novel adaptive genotypes in the face of climate change and other stressors (Habel and Schmitt, 2009; de Jong et al., 2023). Hence, *insulanus* may benefit from management approaches that focus on increasing genetic diversity through genetic rescue and expanding the number of populations through translocation of captive-reared individuals (refer to *Considerations for species management section* below).

Additionally, some individuals had high inbreeding coefficients (Figure S2), indicating there is a risk of inbreeding depression for the species if unconstrained inbreeding continues. Inbreeding depression in butterflies has been shown to reduce thermal tolerance (Dierks et al., 2012a), decrease fertility and larval survival (van Oosterhout et al., 2000; Cassel et al., 2001; Nieminen et al., 2001), decrease adult lifespan (Saccheri et al., 1998; De Nardin et al., 2016), reduce adaptive potential (Dierks et al., 2012b), and ultimately increase the risk of extinction (Saccheri et al., 1998; Nonaka et al., 2019). The negative consequences of inbreeding depression largely result from the unmasking of deleterious alleles, when inbreeding leads to a higher number of individuals expressing recessive alleles with negative fitness consequences (Kyriazis et al., 2021), and additional studies to quantify inbreeding depression in *insulanus* may prove helpful to the conservation of the species. Furthermore, since inbreeding coefficients varied greatly among individuals and pairwise relatedness indicated that most individuals collected for this study were not closely related, careful selection of individuals during reintroduction and/or translocation could help to mitigate the effects of inbreeding.

Genetic diversity is not the only measure to consider when it comes to survival of the species. For example, low heterozygosity and low population size in the Uncompahgre fritillary (*Boloria acrocnema*) led Britten et al. (1994) to predict the species’ extinction as “inevitable.” Instead, the species persisted and maintained genetic diversity over time, rather than succumbing to the effects of genetic erosion (Monroe et al., 2016). Although low genetic diversity is a serious risk factor for extinction, it must be considered within the context of a species’ ecology and demographic history to determine the likely impact (Schultz et al., 2019; Teixeira and Huber, 2021; DiLeo et al., 2024). The next section provides more detail on how these results could be integrated into management considerations to improve the long-term outlook of the species.

## Considerations for species management

*Genetic rescue.* Due to its small effective population size, low levels of genetic variation, and evidence of inbreeding, *insulanus* may be a good candidate for genetic rescue (Fitzpatrick et al., 2023), whereby new alleles are introduced into a population (i.e., gene flow) to ameliorate the deleterious effects of low genetic diversity and decrease the likelihood of extinction (Bell et al., 2019; Hohenlohe et al., 2021). Increasing gene flow by improving connectivity between populations has been successful in slowing the erosion of genetic diversity in other butterfly species, such as the Rocky Mountain parnassian (*Parnassius smintheus*) (Jangjoo et al., 2016), but this strategy is impractical for *insulanus*, which is island-bound and only has one population. Instead, limited hybridization of *insulanus* with one of the closely related subspecies could be an option for increasing genetic diversity. This approach was successfully used with an isolated population of the Marsh Fritillary butterfly (*Euphydryas aurinia*), which benefited from an influx of new alleles but still maintained genetic distinctiveness (Davis et al., 2021). In addition to increasing genetic diversity, genetic rescue can increase population growth rates, which may be necessary to provide enough surplus individuals to translocate into new populations (Bell et al., 2019; Berger-Tal et al., 2020). However, whereas there was only weak divergence between the mitogenomes of *insulanus* and some of the other subspecies (i.e., *ausonides* and *mayi*), a thorough analysis of the differences in nuclear genetic divergence between *insulanus* and the mainland subspecies would provide further information on whether genetic rescue through selective hybridization is feasible and identify potential genetic incompatibilities to reduce the risk of outbreeding depression.

*Establishment of new populations.* Currently, *insulanus* is only found in one small population, leaving it acutely vulnerable to stochastic events. We also identified multiple highly related (i.e., likely full sibling or higher) and inbred individuals which, when reintroduced into the same location after captive rearing, may contribute to an increase in inbreeding depression. One option to consider for mitigating these issues is to translocate highly related individuals to different locations where they could form new populations. Translocation, or movement of individuals from one location to another, can be an important tool for threatened and endangered species (Schultz et al., 2008; Daniels et al., 2018). For example, the large blue (*Maculinea arion*), scarce large blue (*Phengaris teleius*), chequered skipper (*Carterocephalus palaemon*), and clouded Apollo (*Parnassius mnemosyne*) were all successfully translocated and able to establish new populations that persisted >5 years (Andersen et al., 2014; Kuussaari et al., 2015; Bourn et al., 2024; Sánchez-García et al., 2024). The few studies that have tracked genetic diversity in translocated populations have been mixed; whereas *M. arion* showed evidence of new allelic diversity, *P. teleius* lost allelic diversity over time, likely due to founder effects (Andersen et al., 2014; Sánchez-García et al., 2024). Founder effects and existing inbreeding load also likely led to deformed wings in a translocated population of the Apollo butterfly (*Parnassius apollo*) (Witkowski et al., 1997; Adamski and Witkowski, 1999). Consequently, *insulanus* may benefit from additional research to quantify inbreeding load to reduce the likelihood of deleterious phenotypes arising in the new populations created through translocation.

*Introduction of novel larval host plants.* Larval host plant specificity has been identified as a major extinction risk for butterflies (Koh et al., 2004; Palash et al., 2022). Guppy and Shepard (2001) suggested that hairy rock cress (*Arabis eschscholtziana*, previously *A. hirsuta*) may have been the original larval host plant for *insulanus*. In addition to a native pepperweed (*Lepidium virginicum* var. *menziesii*), *insulanus* included in this study were collected from the recently introduced tumble mustard (*Sisymbrium altissimum*) and field mustard (*Brassica rapa*). Lambert (2011) had some success using tower mustard (*Turritis glabra*) as a larval host plant for *insulanus,* providing further evidence of plasticity in host plant use in this species. Other research documenting host plant use by *Euchloe ausonides* subspecies show that they use a variety of weedy mustards (Brassicaceae) (Opler, 1974; Guppy and Shepard, 2001; Back et al., 2011), which could provide additional options for oviposition and larval food sources if made available. This flexibility in host plant use is underscored by our results, which show a lack of population structure and relatedness by host plant, suggesting that *insulanus* females do not exhibit strong natal fidelity when choosing ovipositing sites (Nice et al., 2002; Downey and Nice, 2011).

Since adults lay eggs on unopened flower buds, the species may be susceptible to shifts in flowering phenology due to climate change, which could lead to asynchrony between flower budding and ovipositing (Hill et al., 2021). Maintaining access to a range of hosts may increase the likelihood that flower budding in at least one host species will continue to overlap with the timing of egg deposition (Navarro-Cano et al., 2015). Providing access to additional host plants could therefore prove useful in reducing the vulnerability of *insulanus* to environmental stressors. However, recently introduced non-native host plants can also act as ecological traps when novel phytochemicals negatively impact larval development or lead to phenotypic changes with knock-on effects to reproduction (Braga, 2023; Jarrett and Miller, 2024). Thus, additional studies on the suitability of novel larval host plants for *insulanus* are warranted prior to their introduction to existing habitat or use as larval food during captive rearing.

## Considerations for ongoing genetic monitoring

*Marker choice*. Molecular methods have improved substantially since this study began in 2014, and the adoption of more advanced techniques may be useful for ongoing genetic monitoring. For example, the potential issues of using capillary electrophoresis for genotyping microsatellites (i.e., difficulty transferring between laboratories, subjectivity in scoring, and labor intensiveness have been largely resolved by newer sequence-based microsatellite genotyping methods (such as SSRseq, also known as genotyping-by-sequencing [GBS]) (De Barba et al., 2017). Whereas capillary electrophoresis relies on visually scoring alleles by sequence length, SSRseq involves sequencing the entire microsatellite (Darby et al., 2016; Vartia et al., 2016). Alleles can then be called bioinformatically to produce typical genotypes (Barbian et al., 2018; Tibihika et al., 2019) or the entire sequence can be used to generate high-resolution haplotypes (Lepais et al., 2020), reducing the risk of false alleles being called. SSRseq also increases genotyping success with degraded DNA compared to capillary electrophoresis (De Barba et al., 2017), hence potentially resolving the issues with allele dropout seen with some loci in this study. Nevertheless, not all pre-existing microsatellites perform well with SSRseq, so implementing this method could require additional microsatellites to be developed from the candidate sequences generated for this study.

*Source of genetic material*. Our results indicate that meconium collected on FTA cards, larval exuviae, and pupae from natural mortalities all generally produced high enough quality DNA to be used for microsatellite genotyping, while frass did not. However, there were issues with allele dropout for the microsatellite data, which needed to be accounted for when assessing the results. Limiting genetic monitoring using opportunistically collected samples and natural mortalities could also bias the results, since most population genetic analyses assume that samples are randomly selected and representative of the population (Phillips et al., 2019). These biases may be reflected in the Structure results, which showed more clustering among individuals collected in 2018, possibly due to higher relatedness among individuals from non-random sampling skewing results (Schwartz and McKelvey, 2009; Puechmaille, 2016).

To mitigate these issues, it could be advantageous to collect fresh tissue samples from the general population, rather than incidental samples from larvae in the captive rearing program. Sample design that limits the collection of too many individuals from one area, coupled with methods such as non-lethal wing clipping and/or leg removal, could provide higher quality DNA and reduce bias in the data. Wing clipping and leg removal have been shown to have no significant impact on behavior or survival in multiple species of butterfly (Hamm et al., 2010; Koscinski et al., 2011; Crawford et al., 2013). Wing clipping was previously approved by the USFWS for use in at least two endangered butterfly species (i.e., Mitchell’s satyr, *Neonympha mitchellii mitchellii*, and Karner blue, *Plebejus samuelis*) (Hamm et al., 2014; Zhang et al., 2024), and thus may be an effective technique for obtaining higher quality DNA from adults that could be used to improve the accuracy of microsatellite-based monitoring or provide the material needed to more fine-scale genomic analyses (Webster et al., 2023).

## Conclusion

*Euchloe ausonides insulanus* has lost most of its habitat over the last ∼100 years, leading to a marked reduction in population size. The results on this study underscore those demographic changes, providing genetic evidence that there is likely a single population remaining, with low genetic diversity and a small effective population size. However, there is often a temporal lag between population declines and genetic erosion, leading to an overestimate of genetic diversity measures immediately following demographic changes (Gargiulo et al., 2025). Hence, the full effects of the demographic events that have been affecting may not yet be reflected in the genetic diversity shown in this study and further declines in genetic diversity may continue to occur even if the population remains stable going forward. Given these concerns, conservation strategies such as genetic rescue and managed translocation may be necessary to mitigate inbreeding effects and bolster genetic diversity. The success of such interventions in other butterfly species suggests that similar approaches could increase adaptive potential and improve the long-term viability of *insulanus* in the face of ongoing environmental and demographic challenges.

## Acknowledgments

We thank the following individuals for their contributions to this study: Jing Zhang, Qian Cong, and Nick Grishin (University of Texas Southwestern Medical Center) for providing mitochondrial sequence data and specimen metadata; Michael Eackles (USGS) for assisting with lab work, designing the multiplex PCRs, and microsatellite genotyping; Jenny Shrum (USFWS) for providing samples and metadata; Karen Reagan (USGS) for initial conceptualization of the study and procuring samples; Erin Sullivan (Woodland Park Zoo) for providing samples of additional *E. ausonides* subspecies; San Juan Island National Historical Park (NPS) staff for assisting with sample collection; Jaret Daniels (McGuire Center for Lepidoptera and Biodiversity, Florida Museum of Natural History) for assisting with initial study planning; Julie Combs (Washington Department of Fish and Wildlife) for additional information on the natural and management history of *insulanus*; and Erin Adams (USFWS) and two USGS peer reviewers for their helpful comments and suggestions on a draft of this manuscript. Any use of trade, firm, or product names is for descriptive purposes only and does not imply endorsement by the U.S. Government.

## Statements and Declarations

### Funding

This work was supported by the United States Geological Survey Science Support Program in FY2017-2020 and FY2021-FY2024.

*Competing interests*: The authors have no relevant financial or non-financial interests to disclose.

### Author contributions

All authors contributed to the study conception and design. Material preparation, data collection, and lab work was performed by Aaron Aunins, Colleen Young, and Robin Johnson.

Data analyses were performed and interpreted by Kara Jones, Aaron Aunins, and Cheryl Morrison. The first draft of the manuscript was written by Kara Jones, Aaron Aunins, and Cheryl Morrison. All authors approved the final manuscript.

## Data availability

Sequencing data and associated sample metadata were deposited to the National Center for Biotechnology Information (NCBI) under BioProject PRJNA1156227 (https://doi.org/10.5066/P136HXY4). The *Euchloe ausonides insulanus* annotated mitochondrial genome was deposited on GenBank under accession PQ287242. Microsatellite genotypes, ddRAD single nucleotide polymorphisms (SNPs), and additional sample metadata were deposited on Zenodo (https://doi.org/10.5281/zenodo.14502991). At the time of publication, additional sequences used in the mitogenome phylogeny from the University of Texas Southwestern Medical Center not been made available for publication.

## Notes

https://doi.org/10.5066/P136HXY4

https://doi.org/10.5281/zenodo.14502991

https://www.ncbi.nlm.nih.gov/nuccore/PQ287242.1

https://www.ncbi.nlm.nih.gov/bioproject/PRJNA1156227

### Competing Interest Statement

The authors have declared no competing interest.

