## Supplemental Tables and Figures for "Population genetics of the endangered narrowly endemic Island Marble butterfly (*Euchloe ausonides insulanus*)"

Supplemental Materials for Conservation Genetics and Genomics of the Narrowly Endemic Island Marble Butterfly: Assessing Population Structure, Facilitating Supportive Breeding, and Identifying Adaptive Potential

Kara S. Jones, Aaron W. Aunins, Colleen C. Young, Robin L. Johnson, Cheryl L. Morrison

U.S. Geological Survey, Eastern Ecological Science Center, Leetown Research Laboratory, 11469, Leetown Road, Kearneysville, WV

Any use of trade, firm, or product names is for descriptive purposes only and does not imply endorsement by the U.S. Government.

Table of Contents

**Table S1.** Samples used in this study. All samples were from Euchloe ausonides insulanus, except for EaC3 and EaC8 which were from unidentified Euchloe ausonides subspecies. If the larval instar stage was known, it is noted in parentheses in the “Sample type” column (e.g., I, II, III, IV, V). If the pupal stage is known, it is noted whether the larva never emerged (uneclosed) or only partially emerged (semi-eclosed) from the chrysalis. Refer to Table 1 for further details on sample types.	4

**Table S2.** Primer sets and characteristics of microsatellite loci. PET, VIC, NED, and FAM refer to the names of 5’ reporter dyes available from Applied Biosystems (Waltham, Massachusetts). Microsatellite locus genotypes deposited on Zenodo: https//doi.org/10.5281/zenodo.14502991	12

**Table S3.** Primer pairs for mitogenomic long range PCR. Location refers to the bp position in the mitogenome when oriented to start at the beginning of the COX1 gene. Amplicon size refers to the size of the amplified PCR product for each primer pair. Primers were modified from those originally described by Park et al. (2012). Sequences generated using long range PCR are available under NCBI Project accession PRJNA1156227: https://doi.org/10.5066/P136HXY4	13

**Table S4.** List of the sample metadata for additional mitogenome sequences obtained from an unrelated sequencing project (Zhang et al., University of Texas Southwestern Medical Center, unpublished data, 2023) and used in the mitogenome phylogeny.	14

**Table S5.** Results from initial analysis of microsatellite markers, including number of alleles (N_a_), number of effective alleles (A_e_), observed heterozygosity (H_O_), expected heterozygosity (H_e_), inbreeding coefficient (F), and p-value for Hardy-Weinberg equilibrium (HWE). * significant at an alpha level of 0.05; ** significant at an alpha level of 0.01; *** significant at an alpha level of 0.001. Microsatellite genotypes used in this analysis are available on Zenodo: https://doi.org/10.5066/P136HXY4	16

**Table S6.** Polymorphic information content (PIC), probability of identity (P_(ID)_), and probability of identity for siblings (P_(ID)sib_) for all 13 microsatellite loci. Microsatellite genotypes used in this analysis are available on Zenodo: https://doi.org/10.5066/P136HXY4	19

**Table S*7*.** Full sibship groups assigned by COLONY for each of the three-year cohorts. For analyses where siblings were removed, only one individual was kept from groups with an inclusive probability ≥0.9. Microsatellite genotypes used in this analysis are available on Zenodo: https://doi.org/10.5066/P136HXY4	20

**Table S8.** Results from mitogenome assembly using shotgun sequencing and long range PCR. All libraries used for mitogenome assembly were sequenced with 250 bp paired-end (PE) reads, except for Eai-S43, which used 150 bp PE. Assembly did not result in a circular mitogenome for Eai-162 and EaC8. Depth = mean read depth of mitogenome assembly ± standard deviation. E. a. subsp. = unknown Euchloe ausonides subspecies. Data: https://doi.org/10.5066/P136HXY4	22

**Figure S1.** Estimated error rates for each locus from each year cohort analyzed with COLONY. Starting error rates for allele dropout and false alleles were both 0.01.	23

**Figure S2.** Inbreeding coefficients (F_IS_) calculated by EMIBD9 for each individual, grouped by year. Jitter has been to the points to help distinguish points.	24

**Figure S3.** Comparison of estimated ancestry from Structure analysis for population clusters (K) = 2-4 for each year. Individuals are grouped by larval host plant genus within each year and colors indicate estimated proportion of estimated ancestry to each cluster. Microsatellite genotypes used in this analysis are available on Zenodo: https://doi.org/10.5066/P136HXY4	25

**Figure S4.** Map of mitogenome for Euchloe ausonides insulanus showing the location of 13 protein-coding genes, 22 tRNAs, 2 rRNAs, D-loop, and replication origin (OH), along with GC content. Mitogenomes from all insulanus individuals sequenced were identical. The annotated mitogenome used to create this figure is available under GenBank accession PQ287242.1. Photograph of insulanus courtesy of Karen Reagan/USFWS (Public Domain), 2016.	26

**Table S1.** Samples used in this study. All samples were from *Euchloe ausonides insulanus*, except for EaC3 and EaC8 which were from unidentified *Euchloe ausonides* subspecies. If the larval instar stage was known, it is noted in parentheses in the “Sample type” column (e.g., I, II, III, IV, V). If the pupal stage is known, it is noted whether the larva never emerged (uneclosed) or only partially emerged (semi-eclosed) from the chrysalis. Refer to Table 1 for further details on sample types.

| **Sample ID** | **Sample type** | **Year collected** | **Species** |
| --- | --- | --- | --- |
| 13SI8 | Pupa | 2013 | *E. a. insulanus* |
| 16BR05 | Meconium | 2016 | *E. a. insulanus* |
| 16BR17 | Exuvia | 2017 | *E. a. insulanus* |
| 16BR20 | Pupa (semi-eclosed) | 2017 | *E. a. insulanus* |
| 16BR23 | Exuvia | 2017 | *E. a. insulanus* |
| 16BR28 | Meconium | 2016 | *E. a. insulanus* |
| 16BR63 | Pupa (semi-eclosed) | 2017 | *E. a. insulanus* |
| 16LE04 | Meconium | 2016 | *E. a. insulanus* |
| 16SI04 | Pupa (uneclosed) | 2017 | *E. a. insulanus* |
| 16SI05 | Meconium | 2016 | *E. a. insulanus* |
| 16SI06 | Pupa (uneclosed) | 2017 | *E. a. insulanus* |
| 16SI36 | Meconium | 2016 | *E. a. insulanus* |
| 16SI40 | Meconium | 2016 | *E. a. insulanus* |
| 16SI58 | Meconium | 2016 | *E. a. insulanus* |
| 16SI70 | Meconium | 2016 | *E. a. insulanus* |
| 17BR02 | Meconium | 2018 | *E. a. insulanus* |
| 17BR09 | Meconium | 2018 | *E. a. insulanus* |
| 17BR100 | Meconium | 2018 | *E. a. insulanus* |
| 17BR101 | Meconium | 2018 | *E. a. insulanus* |
| 17BR16 | Pupa | 2017 | *E. a. insulanus* |
| 17BR18 | Meconium | 2018 | *E. a. insulanus* |
| 17BR29 | Meconium | 2018 | *E. a. insulanus* |
| 17BR34 | Meconium | 2018 | *E. a. insulanus* |
| 17BR42 | Meconium | 2018 | *E. a. insulanus* |
| 17BR49 | Meconium | 2018 | *E. a. insulanus* |
| 17BR52 | Meconium | 2018 | *E. a. insulanus* |
| 17BR62 | Larva | 2017 | *E. a. insulanus* |
| 17BR65 | Pupa | 2017 | *E. a. insulanus* |
| 17BR67 | Meconium | 2018 | *E. a. insulanus* |
| 17BR73 | Meconium | 2018 | *E. a. insulanus* |
| 17BR79 | Meconium | 2018 | *E. a. insulanus* |
| 17BR85 | Meconium | 2018 | *E. a. insulanus* |
| 17BR86 | Meconium | 2018 | *E. a. insulanus* |
| 17BR88 | Larva | 2017 | *E. a. insulanus* |
| 17BR91 | Meconium | 2018 | *E. a. insulanus* |
| 17BR96 | Meconium | 2018 | *E. a. insulanus* |
| 17BR97 | Meconium | 2018 | *E. a. insulanus* |
| 17BR99 | Meconium | 2018 | *E. a. insulanus* |
| 17LE03 | Meconium | 2018 | *E. a. insulanus* |
| 17LE07 | Larva | 2017 | *E. a. insulanus* |
| 17LE10 | Meconium | 2018 | *E. a. insulanus* |
| 17LE11 | Meconium | 2018 | *E. a. insulanus* |
| 17LE14 | Meconium | 2018 | *E. a. insulanus* |
| 17LE18 | Meconium | 2018 | *E. a. insulanus* |
| 17LE19 | Meconium | 2018 | *E. a. insulanus* |
| 17LE20 | Larva | 2017 | *E. a. insulanus* |
| 17LE22 | Meconium | 2018 | *E. a. insulanus* |
| 17LE26 | Larva | 2017 | *E. a. insulanus* |
| 17LE31 | Meconium | 2018 | *E. a. insulanus* |
| 17SI16 | Meconium | 2018 | *E. a. insulanus* |
| 17SI21 | Larva | 2017 | *E. a. insulanus* |
| 17SI22 | Meconium | 2018 | *E. a. insulanus* |
| 17SI28 | Meconium | 2018 | *E. a. insulanus* |
| 17SI35 | Meconium | 2018 | *E. a. insulanus* |
| 17SI44 | Meconium | 2018 | *E. a. insulanus* |
| 17SI46 | Meconium | 2018 | *E. a. insulanus* |
| 17SI47 | Meconium | 2018 | *E. a. insulanus* |
| 17SI48 | Meconium | 2018 | *E. a. insulanus* |
| 17SI52 | Meconium | 2018 | *E. a. insulanus* |
| 17SI53 | Meconium | 2018 | *E. a. insulanus* |
| 17SI55 | Meconium | 2018 | *E. a. insulanus* |
| 17SI56 | Meconium | 2018 | *E. a. insulanus* |
| 17SI57 | Meconium | 2018 | *E. a. insulanus* |
| 17SI60 | Meconium | 2018 | *E. a. insulanus* |
| 17SI63 | Meconium | 2018 | *E. a. insulanus* |
| 17SI65 | Meconium | 2018 | *E. a. insulanus* |
| 17SI67 | Meconium | 2018 | *E. a. insulanus* |
| 17SI70 | Meconium | 2018 | *E. a. insulanus* |
| 17SI72 | Meconium | 2018 | *E. a. insulanus* |
| 17SI74 | Meconium | 2018 | *E. a. insulanus* |
| 17SI75 | Meconium | 2018 | *E. a. insulanus* |
| 17SI78 | Meconium | 2018 | *E. a. insulanus* |
| 18BR01 | Meconium | 2019 | *E. a. insulanus* |
| 18BR04 | Meconium | 2019 | *E. a. insulanus* |
| 18BR05 | Meconium | 2019 | *E. a. insulanus* |
| 18BR06 | Meconium | 2019 | *E. a. insulanus* |
| 18BR07 | Larva | 2018 | *E. a. insulanus* |
| 18BR11 | Meconium | 2019 | *E. a. insulanus* |
| 18BR12 | Pupa | 2019 | *E. a. insulanus* |
| 18BR15 | Meconium | 2019 | *E. a. insulanus* |
| 18BR16 | Larva | 2018 | *E. a. insulanus* |
| 18BR18 | Larva | 2018 | *E. a. insulanus* |
| 18BR20 | Larva | 2018 | *E. a. insulanus* |
| 18BR21 | Meconium | 2018 | *E. a. insulanus* |
| 18BR22 | Meconium | 2018 | *E. a. insulanus* |
| 18BR23 | Meconium | 2018 | *E. a. insulanus* |
| 18BR24 | Meconium | 2018 | *E. a. insulanus* |
| 18BR25 | Meconium | 2018 | *E. a. insulanus* |
| 18BR27 | Meconium | 2018 | *E. a. insulanus* |
| 18BR28 | Meconium | 2018 | *E. a. insulanus* |
| 18BR29 | Meconium | 2018 | *E. a. insulanus* |
| 18BR30 | Meconium | 2018 | *E. a. insulanus* |
| 18BR31 | Meconium | 2018 | *E. a. insulanus* |
| 18BR34 | Larva | 2017 | *E. a. insulanus* |
| 18BR35 | Meconium | 2018 | *E. a. insulanus* |
| 18BR36 | Meconium | 2018 | *E. a. insulanus* |
| 18BR38 | Meconium | 2018 | *E. a. insulanus* |
| 18BR39 | Meconium | 2018 | *E. a. insulanus* |
| 18BR40 | Meconium | 2018 | *E. a. insulanus* |
| 18BR42 | Meconium | 2018 | *E. a. insulanus* |
| 18BR44 | Meconium | 2018 | *E. a. insulanus* |
| 18BR45 | Meconium | 2018 | *E. a. insulanus* |
| 18BR46 | Meconium | 2018 | *E. a. insulanus* |
| 18BR48 | Meconium | 2018 | *E. a. insulanus* |
| 18BR50 | Meconium | 2018 | *E. a. insulanus* |
| 18BR51 | Meconium | 2018 | *E. a. insulanus* |
| 18BR52 | Meconium | 2018 | *E. a. insulanus* |
| 18BR53 | Meconium | 2018 | *E. a. insulanus* |
| 18BR54 | Meconium | 2018 | *E. a. insulanus* |
| 18BR57 | Meconium | 2018 | *E. a. insulanus* |
| 18BR58 | Meconium | 2018 | *E. a. insulanus* |
| 18BR59 | Meconium | 2018 | *E. a. insulanus* |
| 18LE01 | Meconium | 2018 | *E. a. insulanus* |
| 18LE05 | Meconium | 2018 | *E. a. insulanus* |
| 18LE06 | Meconium | 2018 | *E. a. insulanus* |
| 18LE07 | Meconium | 2018 | *E. a. insulanus* |
| 18LE08 | Meconium | 2018 | *E. a. insulanus* |
| 18LE09 | Meconium | 2018 | *E. a. insulanus* |
| 18LE11 | Meconium | 2018 | *E. a. insulanus* |
| 18LE13 | Meconium | 2018 | *E. a. insulanus* |
| 18SI01 | Meconium | 2018 | *E. a. insulanus* |
| 18SI02 | Meconium | 2018 | *E. a. insulanus* |
| 18SI04 | Meconium | 2018 | *E. a. insulanus* |
| 18SI05 | Meconium | 2018 | *E. a. insulanus* |
| 18SI07 | Meconium | 2018 | *E. a. insulanus* |
| 18SI08 | Meconium | 2018 | *E. a. insulanus* |
| 18SI10 | Meconium | 2018 | *E. a. insulanus* |
| 18SI100 | Meconium | 2018 | *E. a. insulanus* |
| 18SI101 | Meconium | 2018 | *E. a. insulanus* |
| 18SI102 | Meconium | 2018 | *E. a. insulanus* |
| 18SI103 | Larva | 2018 | *E. a. insulanus* |
| 18SI104 | Meconium | 2018 | *E. a. insulanus* |
| 18SI105 | Meconium | 2018 | *E. a. insulanus* |
| 18SI106 | Meconium | 2018 | *E. a. insulanus* |
| 18SI109 | Meconium | 2018 | *E. a. insulanus* |
| 18SI110 | Meconium | 2018 | *E. a. insulanus* |
| 18SI111 | Meconium | 2018 | *E. a. insulanus* |
| 18SI113 | Meconium | 2018 | *E. a. insulanus* |
| 18SI115 | Meconium | 2018 | *E. a. insulanus* |
| 18SI116 | Meconium | 2018 | *E. a. insulanus* |
| 18SI117 | Meconium | 2018 | *E. a. insulanus* |
| 18SI118 | Meconium | 2018 | *E. a. insulanus* |
| 18SI119 | Meconium | 2018 | *E. a. insulanus* |
| 18SI12 | Meconium | 2018 | *E. a. insulanus* |
| 18SI122 | Meconium | 2018 | *E. a. insulanus* |
| 18SI123 | Meconium | 2018 | *E. a. insulanus* |
| 18SI124 | Meconium | 2018 | *E. a. insulanus* |
| 18SI126 | Meconium | 2018 | *E. a. insulanus* |
| 18SI128 | Meconium | 2018 | *E. a. insulanus* |
| 18SI13 | Meconium | 2018 | *E. a. insulanus* |
| 18SI130 | Meconium | 2018 | *E. a. insulanus* |
| 18SI131 | Larva | 2018 | *E. a. insulanus* |
| 18SI132 | Meconium | 2018 | *E. a. insulanus* |
| 18SI14 | Meconium | 2018 | *E. a. insulanus* |
| 18SI15 | Meconium | 2018 | *E. a. insulanus* |
| 18SI16 | Larva | 2018 | *E. a. insulanus* |
| 18SI17 | Meconium | 2018 | *E. a. insulanus* |
| 18SI18 | Meconium | 2018 | *E. a. insulanus* |
| 18SI19 | Meconium | 2018 | *E. a. insulanus* |
| 18SI23 | Meconium | 2018 | *E. a. insulanus* |
| 18SI24 | Meconium | 2018 | *E. a. insulanus* |
| 18SI25 | Meconium | 2018 | *E. a. insulanus* |
| 18SI26 | Meconium | 2018 | *E. a. insulanus* |
| 18SI27 | Meconium | 2018 | *E. a. insulanus* |
| 18SI28 | Meconium | 2018 | *E. a. insulanus* |
| 18SI30 | Meconium | 2018 | *E. a. insulanus* |
| 18SI31 | Meconium | 2018 | *E. a. insulanus* |
| 18SI33 | Meconium | 2018 | *E. a. insulanus* |
| 18SI34 | Meconium | 2018 | *E. a. insulanus* |
| 18SI36 | Meconium | 2018 | *E. a. insulanus* |
| 18SI37 | Meconium | 2018 | *E. a. insulanus* |
| 18SI38 | Meconium | 2018 | *E. a. insulanus* |
| 18SI40 | Meconium | 2018 | *E. a. insulanus* |
| 18SI41 | Meconium | 2018 | *E. a. insulanus* |
| 18SI45 | Meconium | 2018 | *E. a. insulanus* |
| 18SI46 | Meconium | 2018 | *E. a. insulanus* |
| 18SI48 | Meconium | 2018 | *E. a. insulanus* |
| 18SI51 | Meconium | 2018 | *E. a. insulanus* |
| 18SI54 | Meconium | 2018 | *E. a. insulanus* |
| 18SI56 | Meconium | 2018 | *E. a. insulanus* |
| 18SI57 | Meconium | 2018 | *E. a. insulanus* |
| 18SI59 | Meconium | 2018 | *E. a. insulanus* |
| 18SI60 | Meconium | 2018 | *E. a. insulanus* |
| 18SI61 | Meconium | 2018 | *E. a. insulanus* |
| 18SI62 | Meconium | 2018 | *E. a. insulanus* |
| 18SI63 | Meconium | 2018 | *E. a. insulanus* |
| 18SI64 | Meconium | 2018 | *E. a. insulanus* |
| 18SI65 | Meconium | 2018 | *E. a. insulanus* |
| 18SI66 | Meconium | 2018 | *E. a. insulanus* |
| 18SI68 | Meconium | 2018 | *E. a. insulanus* |
| 18SI74 | Meconium | 2018 | *E. a. insulanus* |
| 18SI75 | Meconium | 2018 | *E. a. insulanus* |
| 18SI77 | Meconium | 2018 | *E. a. insulanus* |
| 18SI78 | Meconium | 2018 | *E. a. insulanus* |
| 18SI79 | Larva | 2018 | *E. a. insulanus* |
| 18SI80 | Meconium | 2018 | *E. a. insulanus* |
| 18SI81 | Meconium | 2018 | *E. a. insulanus* |
| 18SI82 | Meconium | 2018 | *E. a. insulanus* |
| 18SI83 | Meconium | 2018 | *E. a. insulanus* |
| 18SI84 | Meconium | 2018 | *E. a. insulanus* |
| 18SI85 | Meconium | 2018 | *E. a. insulanus* |
| 18SI86 | Meconium | 2018 | *E. a. insulanus* |
| 18SI87 | Meconium | 2018 | *E. a. insulanus* |
| 18SI88 | Meconium | 2018 | *E. a. insulanus* |
| 18SI90 | Meconium | 2018 | *E. a. insulanus* |
| 18SI92 | Meconium | 2018 | *E. a. insulanus* |
| 18SI94 | Meconium | 2018 | *E. a. insulanus* |
| 18SI95 | Meconium | 2018 | *E. a. insulanus* |
| 18SI96 | Meconium | 2018 | *E. a. insulanus* |
| 19LE007 | Egg | 2019 | *E. a. insulanus* |
| 19LE008 | Larva (I) | 2019 | *E. a. insulanus* |
| 19LE011 | Egg | 2019 | *E. a. insulanus* |
| 19LE017 | Larva (V) | 2019 | *E. a. insulanus* |
| 19LE019 | Larva (I) | 2019 | *E. a. insulanus* |
| 19LE020 | Pupa | 2019 | *E. a. insulanus* |
| 19SI014 | Egg | 2019 | *E. a. insulanus* |
| 19SI016 | Larva (III) | 2019 | *E. a. insulanus* |
| 19SI019 | Larva (V) | 2019 | *E. a. insulanus* |
| 19SI020 | Egg | 2019 | *E. a. insulanus* |
| 19SI021 | Egg | 2019 | *E. a. insulanus* |
| 19SI030 | Egg | 2019 | *E. a. insulanus* |
| 19SI031 | Egg | 2019 | *E. a. insulanus* |
| 19SI039 | Egg | 2019 | *E. a. insulanus* |
| 19SI040 | Egg | 2019 | *E. a. insulanus* |
| 19SI049 | Larva (II) | 2019 | *E. a. insulanus* |
| 19SI063 | Larva (I) | 2019 | *E. a. insulanus* |
| 19SI077 | Larva (I) | 2019 | *E. a. insulanus* |
| 19SI081 | Larva (I) | 2019 | *E. a. insulanus* |
| 19SI082 | Egg | 2019 | *E. a. insulanus* |
| 19SI084 | Egg | 2019 | *E. a. insulanus* |
| 19SI100 | Egg | 2019 | *E. a. insulanus* |
| 19SI111 | Pupa | 2019 | *E. a. insulanus* |
| 19SI116 | Larva (V) | 2019 | *E. a. insulanus* |
| 19SI117 | Larva (I) | 2019 | *E. a. insulanus* |
| 19SI124 | Larva (III) | 2019 | *E. a. insulanus* |
| 19SI125 | Egg | 2019 | *E. a. insulanus* |
| 19SI126 | Larva (I) | 2019 | *E. a. insulanus* |
| 19SI132 | Egg | 2019 | *E. a. insulanus* |
| 19SI142 | Larva (I) | 2019 | *E. a. insulanus* |
| 19SI150 | Larva (V) | 2019 | *E. a. insulanus* |
| 19SI153 | Larva (II) | 2019 | *E. a. insulanus* |
| 19SI155 | Larva (V) | 2019 | *E. a. insulanus* |
| 19SINA | Larva (V) | 2019 | *E. a. insulanus* |
| Eai20-FTA-013 | Meconium | 2019 | *E. a. insulanus* |
| Eai20-FTA-014 | Meconium | 2019 | *E. a. insulanus* |
| Eai20-FTA-015 | Meconium | 2019 | *E. a. insulanus* |
| Eai20-FTA-016 | Meconium | 2019 | *E. a. insulanus* |
| Eai20-FTA-017 | Meconium | 2019 | *E. a. insulanus* |
| EaC3 | Whole organism | 2017 | *E. a. subsp.* |
| EaC8 | Whole organism | 2017 | *E. a. subsp.* |
| Eai-163 | Larva | 2017 | *E. a. insulanus* |
| Eai-79 | Pupa (semi-eclosed) | 2017 | *E. a. insulanus* |
| Eai-S34 | Meconium | 2014 | *E. a. insulanus* |
| Eai20-FTA-150 | Meconium | 2019 | *E. a. insulanus* |
| Eai20-FTA-151 | Meconium | 2019 | *E. a. insulanus* |
| Eai20-FTA-152 | Meconium | 2019 | *E. a. insulanus* |
| Eai20-FTA-153 | Meconium | 2019 | *E. a. insulanus* |
| Eai20-FTA-155 | Meconium | 2019 | *E. a. insulanus* |

**Table S2.** Primer sets and characteristics of microsatellite loci. PET, VIC, NED, and FAM refer to the names of 5’ reporter dyes available from Applied Biosystems (Waltham, Massachusetts). Microsatellite locus genotypes deposited on Zenodo: https//doi.org/10.5281/zenodo.14502991

| **Locus** | **Primer Sequence** | **Size (bp)** | **Repeat motif** | **Multiplex** | **Dye** |
| --- | --- | --- | --- | --- | --- |
| **Locus-11** | F: CAATCGCTCCCACACATCCT  R: TGGTGCAAGAATCCTGGGAA | 111-114 | (AAT)^6^ | 1 | PET |
| **Locus-24** | F: CTCTCCTGTATATGTGTAGACTTTAGA  R: GTGGACACGGTGATGCTAGT | 137-170 | (ACT)^7^ | 1 | VIC |
| **Locus-71** | F: AGCAGTTGGTTGGTCTTCATCA  R: AGGCAAAGTGATTGATAACAATGCA | 173-185 | (ACAT)^6^ | 1 | NED |
| **Locus-40** | F: ACCATGAAATACACCTACGACAG  R: CCATCGCATTGGTCTTAGGC | 144-147 | (ATC)^6^ | 2 | PET |
| **Locus-54** | F: TGGTCAAAGTACGAGTATAACATTCT  R: TTGTGTGGATTTGCAGTGCA | 171-174 | (AAGT)^5^ | 2 | NED |
| **Locus-05** | F: TGTATTGGTTTGGCAGGATAAAG  R: ACAAGCACGTAGGCTTTGTT | 95-98 | (AAC)^6^ | 3 | NED |
| **Locus-58** | F: TCGGATGGATATCTCACTCCT  R: AGATTAATTCACTCATAGATGCATGTT | 134-160 | (AATG)^5^ | 3 | VIC |
| **Locus-60** | F: AGTCTACGTGTAGTATCGAGTAAGA  R: ATTTCTGTGTGTTTGGATGCTAAA | 145-153 | (AATG)^5^ | 3 | FAM |
| **Locus-65** | F: TGGTTAAGCCTCAAAGAACTGA  R: CTCAGACCAATTATGACTAGGTCA | 98-106 | (AATG)^5^ | 3 | PET |
| **Locus-14** | F: TAGCTACGAACCATGTGGGC  R: TGGGCAATTATCGGATTTAAATCA | 120-126 | (AAT)^8^ | 4 | VIC |
| **Locus-20** | F: TCGTCAAACACTGGACCGAA  R: ACAGGCATAATTCTAAGAAGCGC | 181-190 | (AAT)^8^ | 4 | PET |
| **Locus-33** | F: ACATTGTACGCAGAGCCTGT  R: ACATTGGTTCTTTAAAGACTTACCGT | 158-173 | (ATC)^6^ | 4 | NED |
| **Locus-68** | F: ACACAAGTAATCCAGTGGACACT  R: TCGCATTAATCGATAGGGTAAATCG | 118-166 | (AATT)^6^ | 4 | FAM |

**Table S3.** Primer pairs for mitogenomic long range PCR. Location refers to the bp position in the mitogenome when oriented to start at the beginning of the COX1 gene. Amplicon size refers to the size of the amplified PCR product for each primer pair. Primers were modified from those originally described by Park et al. (2012). Sequences generated using long range PCR are available under NCBI Project accession PRJNA1156227: <https://doi.org/10.5066/P136HXY4>

| **Primer pair** | **Sequence** | **Location (bp)** | **Annealing temp (C)** | **Amplicon size (bp)** |
| --- | --- | --- | --- | --- |
| Lep-COI-F1 | CTCTACTAATCATAAAGATATTGG | 19-4086 | 53 | ~4067 |
| LF01-S06-R2 | GATTGGAAGTCAAATATACT |  |  |  |
| LF01-S05-F2 | TTTTGTTTAATAATTTTTTAGG | 2816-7097 | 51 | ~4281 |
| LepND4-R1 | ATTGGTCATGGATTATGTTCATC |  |  |  |
| Lep-ND5-F1 | CTAAAAGGAATTTGAGCTCT | 6019-11788 | 56 | ~5769 |
| Lep-lrRNA-R1 | CTGTACAAAGGTAGCATAATAAAT |  |  |  |
| Lep-lrRNA-F1 | TGTAAGATTTTAATGATCGAACAGAT | 11340-721 | 58 | ~4575 |
| LepCOI-R1 | CTTCAGGATGTCCGAAAAATC |  |  |  |

**Table S4.** List of the sample metadata for additional mitogenome sequences obtained from an unrelated sequencing project (Zhang et al., University of Texas Southwestern Medical Center, unpublished data, 2023^[1]^) and used in the mitogenome phylogeny.

| **Sample ID** | **Taxon name** | **State** | **County/Region** | **Date collected** |
| --- | --- | --- | --- | --- |
| 6465 | *E. a. coloradensis* | Colorado | Grand Co. | 6-Jul-2016 |
| 8855 | *E. a. coloradensis* | New Mexico | Santa Fe Co. | 15-May-2017 |
| 10925 | *Euchloe lotta* | Arizona | Pima Co. | 7-Apr-2018 |
| NVG-01506A05 | *E. a. insulanus* | British Columbia | Unknown | 3-Jun-2004 |
| NVG-01506A05 | *E. a. insulanus* | British Columbia | Vancouver Island | 3-Jun-2004 |
| NVG-18086H02 | *E. a. ausonides* | California | San Francisco Co.? | Unknown |
| NVG-18086H03 | *E. a. ausonides* | California | San Francisco Co.? | 1851 |
| NVG-19046H04 | *E. a. coloradensis* | Colorado | Jefferson Co. | prior to 1881 |
| NVG-19046H10 | *E. a. palaeoreios* | South Dakota | Lawrence Co. | prior to 1976 |
| NVG-19061D02 | *E. a. ausonides* | California | Santa Clara Co. | 23-May-2005 |
| NVG-19061D04 | *E. a. ausonides* | California | Fremont Co. | Apr-2019 |
| NVG-20039G07 | *E. a. coloradensis* | Utah | Unknown | 2016-2020 |
| NVG-20059E09 | *E. a. coloradensis* | Utah | Summit Co. | 2010 |
| NVG-20067F02 | *E. a. mayi* | Manitoba | Division No. 17 | 24-Jul-1967 |
| NVG-20067F03 | *E. a. mayi* | Manitoba | Division No. 17 | 24-Jul-1967 |
| NVG-20067F04 | *E. a. transmontana* | California | Tulare Co. | 22-May-2012 |
| NVG-20067F05 | *E. a. transmontana* | Nevada | Elko Co. | 10-Jul-1982 |
| NVG-20067F07 | *E. a. transmontana* | Oregon | Jackson Co. | 22-Jun-2014 |
| NVG-20067F08 | *E. a. transmontana* | Washington | Yakima Co. | May-1963 |
| NVG-20067F11 | *E. a. palaeoreios* | South Dakota | Lawrence Co. | 17-Jun-1967 |
| NVG-20067F12 | *E. a. palaeoreios* | South Dakota | Pennington Co. | 12-Jul-1967 |
| NVG-21021D04 | *E. a. transmontana* | California | Tulare Co. | 28-May-2007 |
| NVG-21036E11 | *E. a. transmontana* | Nevada | Lander Co. | 4-Mar-1970 |
| NVG-21036E12 | *E. a. transmontana* | Nevada | Lander Co. | 4-Mar-1970 |
| NVG-21058G04 | *E. a. insulanus* | British Columbia | Vancouver Island | 25-May-1899 |
| NVG-21058G05 | *E. a. insulanus* | British Columbia | Vancouver Island | 17-May-1898 |
| NVG-21058G06 | *E. a. insulanus* | British Columbia | Vancouver Island | 27-May-1898 |
| NVG-21058G07 | *E. a. insulanus* | British Columbia | Vancouver Island | 17-May-1898 |
| NVG-21059B11 | *E. a. ausonides* | California | Marin Co. | 19-May-1960 |
| NVG-21059B12 | *E. a. ausonides* | California | Contra Costa Co. | 5-Jul-2012 |
| NVG-22059E11 | *E. a. ogilvia* | Alaska | North Slope Borough | 14-Jul-2021 |
| PAO1 | *E. a. coloradensis* | Utah | Cache-Utah Co. line | 8-Jun-2016 |
| PAO1004 | *E. a. transmontana* | Colorado | Alpine Co. | Unknown |

**Table S5.** Results from initial analysis of microsatellite markers, including number of alleles (N_a_), number of effective alleles (A_e_), observed heterozygosity (H_O_), expected heterozygosity (H_e_), inbreeding coefficient (F), and p-value for Hardy-Weinberg equilibrium (HWE). * significant at an alpha level of 0.05; ** significant at an alpha level of 0.01; *** significant at an alpha level of 0.001. Microsatellite genotypes used in this analysis are available on Zenodo: https://doi.org/10.5066/P136HXY4

| ***Locus*** | **Population** | **2016** | **2017** | **2018** | **2019** |
| --- | --- | --- | --- | --- | --- |
| ***Eai-05*** | ***n*** | **14** | **56** | **142** | **43** |
|  | *N*_a_ | 2 | 2 | 2 | 2 |
|  | *A*_e_ | 2.000 | 1.994 | 1.969 | 1.996 |
|  | *H*_o_ | 0.143 | 0.482 | 0.551 | 0.442 |
|  | *H*_e_ | 0.500 | 0.499 | 0.492 | 0.499 |
|  | *F* | 0.714 | 0.033 | 0.160 | 0.114 |
|  | *HWE P-val* | 0.008** | 0.805 | 0.50 | 0.453 |
| ***Eai-11*** | ***n*** | **14** | **56** | **137** | **44** |
|  | *N*_a_ | 2 | 2 | 2 | 2 |
|  | *A*_e_ | 1.960 | 1.950 | 1.945 | 1.991 |
|  | *H*_o_ | 0.571 | 0.446 | 0.526 | 0.568 |
|  | *H*_e_ | 0.490 | 0.487 | 0.486 | 0.498 |
|  | *F* | -0.067 | 0.083 | 0.340 | -0.142 |
|  | *HWE P-val* | 0.533 | 0.532 | 0.46 | 0.347 |
| ***Eai-14*** | ***n*** | **13** | **55** | **134** | 44 |
|  | *N*_a_ | 3 | 3 | 3 | 3 |
|  | *A*_e_ | 2.620 | 2.113 | 2.708 | 2.529 |
|  | *H*_o_ | 0.615 | 0.436 | 0.463 | 0.568 |
|  | *H*_e_ | 0.618 | 0.527 | 0.631 | 0.605 |
|  | *F* | 0.005 | 0.172 | 0.266 | 0.060 |
|  | *HWE P-val* | 0.762 | 0.380 | 0.000*** | 0.762 |
| ***Eai-20*** | ***n*** | **12** | **50** | **132** | **42** |
|  | *N*_a_ | 2 | 2 | 3 | 3 |
|  | *A*_e_ | 1.946 | 1.937 | 1.942 | 1.797 |
|  | *H*_o_ | 0.500 | 0.300 | 0.098 | 0.405 |
|  | *H*_e_ | 0.486 | 0.484 | 0.485 | 0.444 |
|  | *F* | -0.029 | 0.380 | 0.797 | 0.088 |
|  | *HWE P-val* | 0.921 | 0.007** | 0.000*** | 0.830 |
| ***Eai-24*** | ***n*** | **13** | **56** | **135** | **44** |
|  | *N*_a_ | 3 | 6 | 7 | 4 |
|  | *A*_e_ | 2.432 | 2.379 | 2.502 | 2.211 |
|  | *H*_o_ | 0.462 | 0.571 | 0.548 | 0.659 |
|  | *H*_e_ | 0.589 | 0.580 | 0.600 | 0.548 |
|  | *F* | 0.216 | 0.014 | 0.087 | -0.203 |
|  | *HWE P-val* | 0.586 | 0.000*** | 0.000*** | 0.077 |
| ***Eai-33*** | ***n*** | **14** | **56** | **135** | **44** |
|  | *N*_a_ | 3 | 2 | 2 | 2 |
|  | *A*_e_ | 2.142 | 1.969 | 1.846 | 1.984 |
|  | *H*_o_ | 0.357 | 0.304 | 0.237 | 0.455 |
|  | *H*_e_ | 0.533 | 0.492 | 0.458 | 0.496 |
|  | *F* | 0.330 | 0.383 | 0.483 | 0.083 |
|  | *HWE P-val* | 0.382 | 0.004** | 0.000*** | 0.580 |
| ***Eai-40*** | ***n*** | **13** | **55** | **137** | **44** |
|  | *N*_a_ | 2 | 2 | 2 | 2 |
|  | *A*_e_ | 1.352 | 1.424 | 1.369 | 1.629 |
|  | *H*_o_ | 0.308 | 0.291 | 0.175 | 0.386 |
|  | *H*_e_ | 0.260 | 0.298 | 0.270 | 0.386 |
|  | *F* | -0.182 | 0.022 | 0.350 | -0.001 |
|  | *HWE P-val* | 0.512 | 0.869 | 0.000*** | 0.996 |
| ***Eai-54*** | ***n*** | **12** | **55** | **136** | **44** |
|  | *N*_a_ | 2 | 2 | 2 | 2 |
|  | *A*_e_ | 1.180 | 1.331 | 1.543 | 1.658 |
|  | *H*_o_ | 0.167 | 0.218 | 0.324 | 0.500 |
|  | *H*_e_ | 0.153 | 0.249 | 0.352 | 0.397 |
|  | *F* | -0.091 | 0.122 | 0.081 | -0.260 |
|  | *HWE P-val* | 0.753 | 0.364 | 0.346 | 0.084 |
| ***Eai-58*** | ***n*** | **14** | **56** | **132** | **44** |
|  | *N*_a_ | 2 | 2 | 3 | 2 |
|  | *A*_e_ | 1.912 | 1.623 | 1.172 | 1.225 |
|  | *H*_o_ | 0.643 | 0.411 | 0.144 | 0.205 |
|  | *H*_e_ | 0.477 | 0.384 | 0.147 | 0.184 |
|  | *F* | -0.348 | -0.070 | 0.021 | -0.114 |
|  | *HWE P-val* | 0.193 | 0.599 | 0.982 | 0.450 |
| ***Eai-60*** | ***n*** | **13** | **56** | **134** | **44** |
|  | *N*_a_ | 2 | 2 | 2 | 2 |
|  | *A*_e_ | 1.352 | 1.392 | 1.312 | 1.337 |
|  | *H*_o_ | 0.308 | 0.304 | 0.187 | 0.250 |
|  | *H*_e_ | 0.260 | 0.282 | 0.238 | 0.252 |
|  | *F* | -0.182 | -0.078 | 0.216 | 0.007 |
|  | *HWE P-val* | 0.512 | 0.562 | 0.012* | 0.962 |
| ***Eai-65*** | ***n*** | **14** | **56** | **135** | **44** |
|  | *N*_a_ | 2 | 2 | 2 | 2 |
|  | *A*_e_ | 1.415 | 1.554 | 1.912 | 1.686 |
|  | *H*_o_ | 0.214 | 0.214 | 0.281 | 0.295 |
|  | *H*_e_ | 0.293 | 0.357 | 0.477 | 0.407 |
|  | *F* | 0.270 | 0.399 | 0.410 | 0.274 |
|  | *HWE P-val* | 0.313 | 0.003** | 0.000*** | 0.069 |
| ***Eai-68*** | ***n*** | **13** | **51** | **135** | **43** |
|  | *N*_a_ | 2 | 2 | 2 | 2 |
|  | *A*_e_ | 1.742 | 1.940 | 2.000 | 1.999 |
|  | *H*_o_ | 0.154 | 0.627 | 0.467 | 0.698 |
|  | *H*_e_ | 0.426 | 0.484 | 0.500 | 0.500 |
|  | *F* | 0.639 | -0.295 | 0.066 | -0.396 |
|  | *HWE P-val* | 0.021* | 0.035* | 0.440 | 0.009** |
| ***Eai-71*** | ***n*** | **14** | **55** | **135** | **43** |
|  | *N*_a_ | 1 | 2 | 3 | 1 |
|  | *A*_e_ | 1.00 | 1.115 | 1.085 | 0 |
|  | *H*_o_ | 0 | 0.109 | 0.037 | 0 |
|  | *H*_e_ | 0 | 0.103 | 0.079 | 0 |
|  | *F* | NA | -0.058 | 0.529 | NA |
|  | *HWE P-val* | NA | 0.669 | 0.000*** | NA |

**Table S6.** Polymorphic information content (PIC), probability of identity (P_(ID)_), and probability of identity for siblings (P_(ID)sib_) for all 13 microsatellite loci. Microsatellite genotypes used in this analysis are available on Zenodo: https://doi.org/10.5066/P136HXY4

|  | **PIC** | **P_(ID)_** | **P_(ID)sib_** |
| --- | --- | --- | --- |
| Locus-05 | 0.37 | 0.38 | 0.60 |
| Locus-11 | 0.37 | 0.38 | 0.60 |
| Locus-14 | 0.54 | 0.23 | 0.50 |
| Locus-20 | 0.37 | 0.38 | 0.60 |
| Locus-24 | 0.67 | 0.13 | 0.43 |
| Locus-33 | 0.37 | 0.38 | 0.60 |
| Locus-40 | 0.25 | 0.54 | 0.73 |
| Locus-54 | 0.28 | 0.50 | 0.71 |
| Locus-58 | 0.21 | 0.60 | 0.78 |
| Locus-60 | 0.22 | 0.59 | 0.77 |
| Locus-65 | 0.34 | 0.41 | 0.63 |
| Locus-68 | 0.37 | 0.38 | 0.60 |
| Locus-71 | 0.06 | 0.87 | 0.93 |
| **Combined** | **0.34** | **0.00001** | **0.00305** |

**Table S*7*.** Full sibship groups assigned by COLONY for each of the three-year cohorts. For analyses where siblings were removed, only one individual was kept from groups with an inclusive probability ≥0.9. Microsatellite genotypes used in this analysis are available on Zenodo: https://doi.org/10.5066/P136HXY4

| **Probability** | **Full siblings** |
| --- | --- |
| **2016-2017** |  |
| 0.9191 | 17BR02,17BR34,17BR49,17SI16,17SI63 |
| 0.9827 | 17BR09,17BR18,17LE07 |
| 0.6335 | 17BR100,17LE14,17SI47,17SI53,17SI56 |
| 0.9907 | 17BR101,17BR65,17SI28,17SI74 |
| 0.7576 | 17BR16,17LE10,17SI72 |
| 0.9613 | 17BR29,17LE18 |
| 0.9706 | 17BR42,17SI35 |
| 1 | 17BR52,17LE31,17SI65 |
| 0.9942 | 17BR62,17SI55 |
| 0.3479 | 17BR67,17SI48 |
| 0.9626 | 17BR73,17BR85,17BR91,17LE03,17SI57,17SI67 |
| 0.8277 | 17BR79,17BR86 |
| 0.8742 | 17BR88,17BR97,17BR99 |
| 1 | 17BR96,17LE26 |
| 0.941 | 17LE11,17SI22 |
| 0.6906 | 17LE19,17SI78 |
| 0.9952 | 17LE20,17SI75 |
| 0.5468 | 17LE22,17SI46 |
| 0.3145 | 17SI52,17SI60,17SI70 |
| **2017-2018** |  |
| 0.9822 | 18BR01,18BR05,18BR06,18BR27,18BR29,18BR44,18SI60 |
| 1 | 18BR04,18BR30,18LE05,18SI12,18SI26,18SI33,18SI74 |
| 1 | 18BR07,18SI16 |
| 0.316 | 18BR11,18BR15,18BR25,18LE07,18SI18,18SI45,18SI80,18SI95 |
| 0.9366 | 18BR12,18LE06,18LE13 |
| 0.9726 | 18BR16,18BR20 |
| 1 | 18BR18,18SI131 |
| 0.7297 | 18BR21,18SI34,18SI40,18SI66,18SI90 |
| 0.4941 | 18BR22,18BR23,18SI04,18SI25,18SI48,18SI59 |
| 1 | 18BR24,18BR48,18BR54,18SI92 |
| 0.2149 | 18BR28,18BR58,18LE01,18SI56,18SI75 |
| 1 | 18BR31,18BR35,18LE09,18LE11 |
| 1 | 18BR34,18SI103 |
| 0.8832 | 18BR36,18BR42,18SI08,18SI28 |
| 0.7356 | 18BR38,18BR40,18SI104,18SI65,18SI83 |
| 0.3714 | 18BR39,18BR51,18SI126,18SI24,18SI46 |
| 0.4681 | 18BR45,18BR46,18SI10,18SI41,18SI63 |
| 0.8411 | 18BR50,18BR52,18BR53 |
| 0.9752 | 18BR57,18SI36,18SI78,18SI81,18SI85 |
| 0.2763 | 18BR59,18SI05,18SI105,18SI109,18SI111,18SI113,18SI115,18SI117,18SI122,18SI123,18SI17 |
| 0.5677 | 18LE08,18SI54,18SI61,18SI64,18SI79 |
| 0.3814 | 18SI01,18SI100,18SI110,18SI132,18SI15,18SI62 |
| 0.3534 | 18SI02,18SI07,18SI101,18SI106,18SI118,18SI19,18SI27,18SI68,18SI96 |
| 0.4605 | 18SI102,18SI31,18SI38,18SI51,18SI84,18SI86,18SI94 |
| 0.9681 | 18SI116,18SI119,18SI124,18SI128,18SI130,18SI14 |
| 1 | 18SI13,18SI57 |
| 0.5303 | 18SI23,18SI37,18SI77,18SI88 |
| 0.5198 | 18SI30,18SI82,18SI87 |
| **2018-2019** |  |
| 0.1553 | 19LE007,19LE017,19LE019,AR02 |
| 0.4092 | 19LE008,19SI100,19SI111,19SI117,AR01,AR06 |
| 0.3097 | 19LE011,19SI150 |
| 0.3079 | 19LE020,19SI084, Eai20-FTA-151,Eai20-FTA-153 |
| 0.403 | 19SI014,19SI016,19SINA, AR04 |
| 0.9984 | 19SI019,19SI132 |
| 0.9676 | 19SI020,19SI039 |
| 0.9974 | 19SI021,19SI126 |
| 0.2188 | 19SI030,19SI031,19SI040,19SI153 |
| 0.3221 | 19SI049,19SI063,19SI142 |
| 1 | 19SI077,19SI082 |
| 0.9947 | 19SI081,19SI124,19SI125 |
| 0.7205 | 19SI116,19SI155 |
| 0.3735 | Eai20-FTA-150, Eai20-FTA-152, Eai20-FTA-155 |

**Table S8.** Results from mitogenome assembly using shotgun sequencing and long range PCR. All libraries used for mitogenome assembly were sequenced with 250 bp paired-end (PE) reads, except for Eai-S43, which used 150 bp PE. Assembly did not result in a circular mitogenome for Eai-162 and EaC8. Depth = mean read depth of mitogenome assembly ± standard deviation. E. a. subsp. = unknown Euchloe ausonides subspecies. Data: https://doi.org/10.5066/P136HXY4

| **Sample** | **Species** | **Raw reads** | **Mitogenome Depth** |
| --- | --- | --- | --- |
| **Shotgun sequencing** | |  |  |
| EaC3 | *E. a. subsp.* | 3,827,208 | 121 ± 141 |
| EaC8 | *E. a. susbp.* | 405,416 | *NA* |
| Eai-S43 | *E. a. insulanus* | 94,216,390 | 49,449 ± 42,720 |
| Eai-160 | *E. a. insulanus* | 3,840,468 | 150 ± 137 |
| Eai-162 | *E. a. insulanus* | 5,835,912 | *NA* |
| Eai-163 | *E. a. insulanus* | 2,342,828 | 63 ± 42.3 |
| Eai-168 | *E. a. insulanus* | 3,578,448 | 162 ± 113 |
| Eai-170 | *E. a. insulanus* | 2,888,624 | 100 ± 91.9 |
| Eai-179 | *E. a. insulanus* | 2,604,518 | 190 ± 84.6 |
| **Long Range PCR** | |  |  |
| EaC3 | *E. a. subsp.* | 1,244,848 | 17,395 ± 12,468 |
| Eai-160 | *E. a. insulanus* | 1,550,916 | 23,195 ± 16,592 |
| Eai-170 | *E. a. insulanus* | 1,157,416 | 16,959 ± 12,980 |
| Eai-179 | *E. a. insulanus* | 747,140 | 10,833 ± 8580.5 |

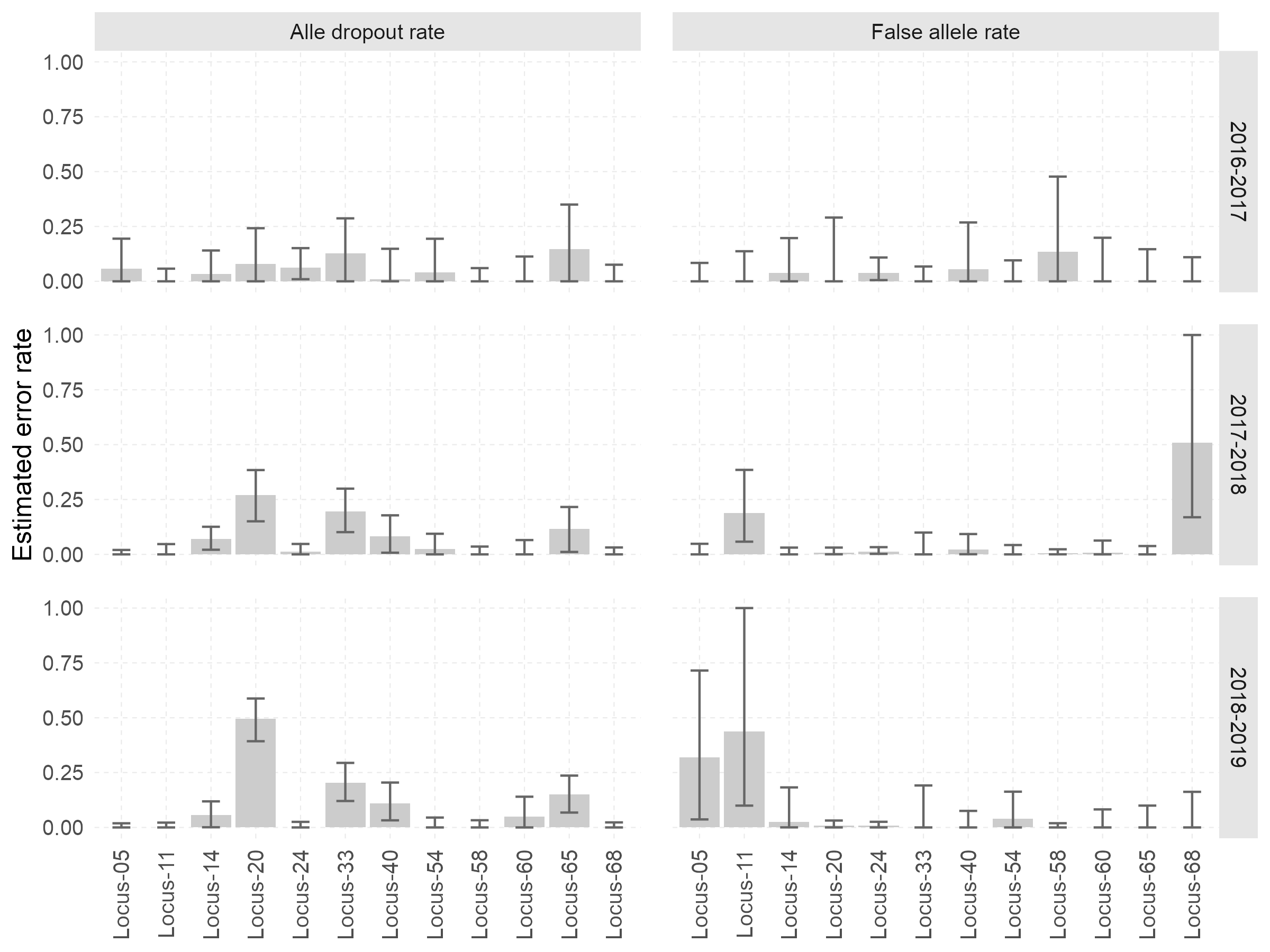

**Figure S1.** Estimated error rates for each locus from each year cohort analyzed with COLONY. Starting error rates for allele dropout and false alleles were both 0.01.

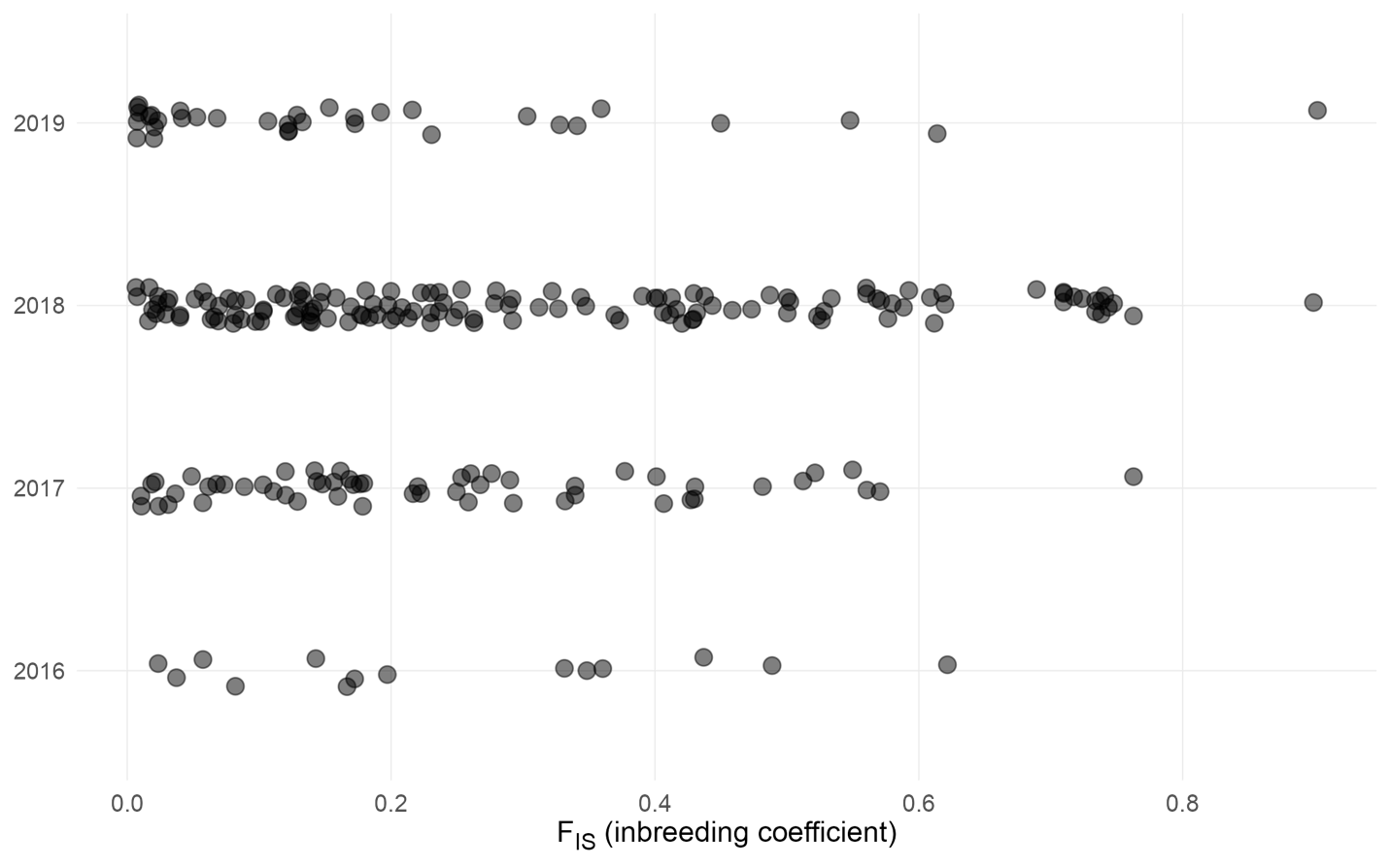

**Figure S2.** Inbreeding coefficients (F_IS_) calculated by EMIBD9 for each individual, grouped by year. Jitter has been to the points to help distinguish points.

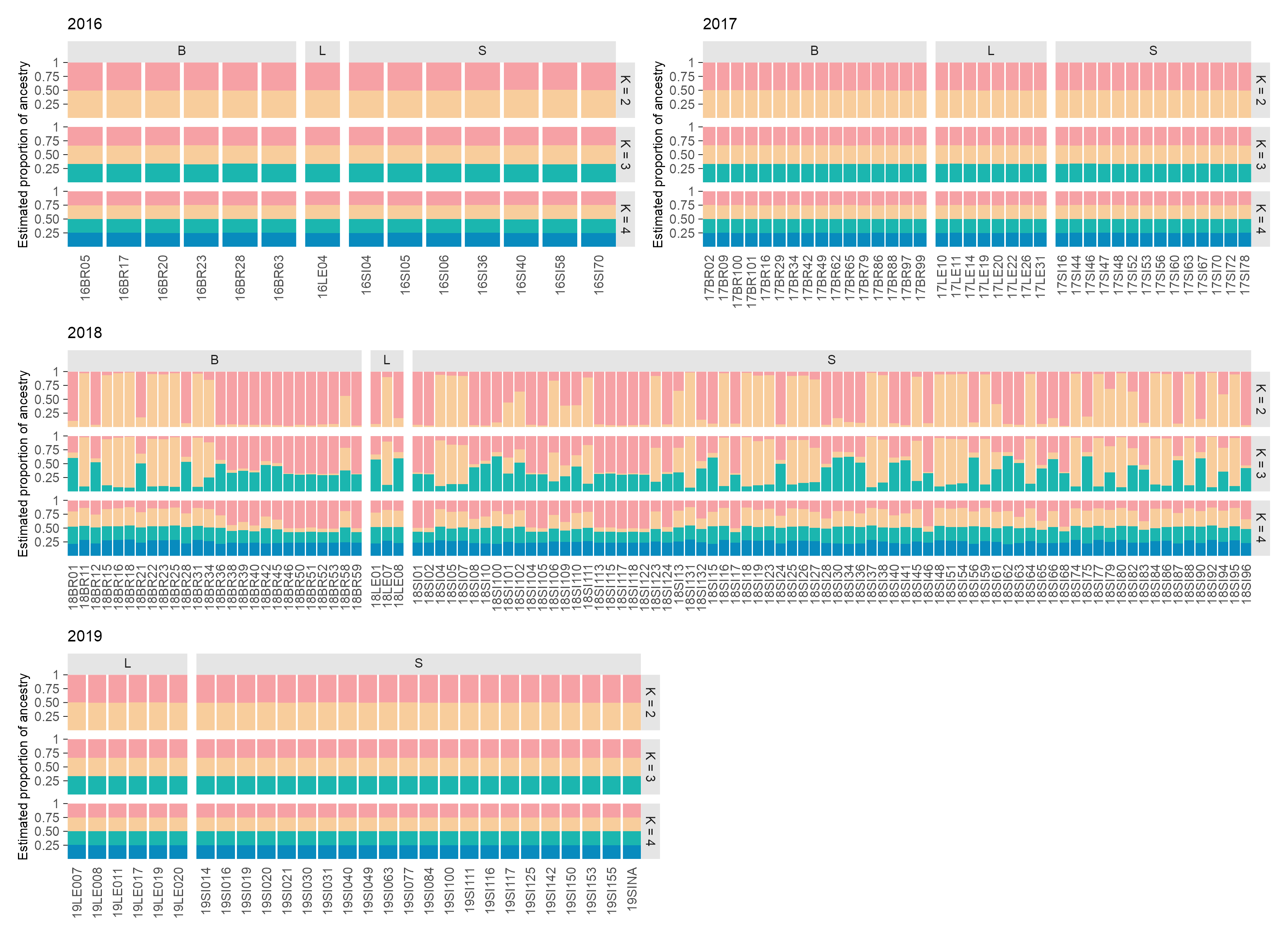

**Figure S3.** Comparison of estimated ancestry from Structure analysis for population clusters (K) = 2-4 for each year. Individuals are grouped by larval host plant genus within each year and colors indicate estimated proportion of estimated ancestry to each cluster. Microsatellite genotypes used in this analysis are available on Zenodo: <https://doi.org/10.5066/P136HXY4>

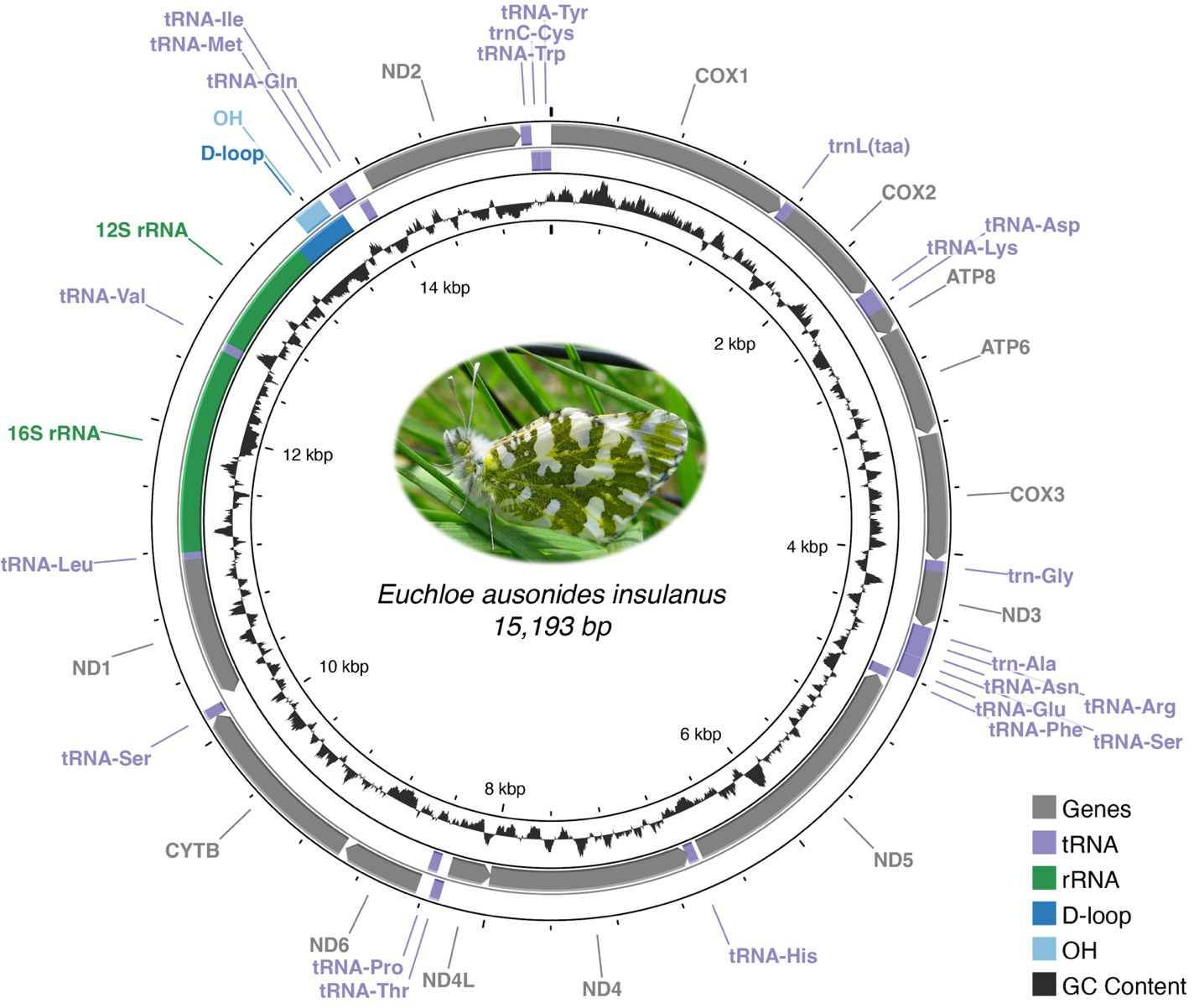

**Figure S4.** Map of mitogenome for Euchloe ausonides insulanus showing the location of 13 protein-coding genes, 22 tRNAs, 2 rRNAs, D-loop, and replication origin (OH), along with GC content. Mitogenomes from all insulanus individuals sequenced were identical. The annotated mitogenome used to create this figure is available under GenBank accession PQ287242.1. Photograph of insulanus courtesy of Karen Reagan/USFWS (Public Domain), 2016.

1. At the time of publication, additional sequences used in the mitogenome phylogeny from the University of Texas Southwestern Medical Center were had not been made available for publication.
